# Epigenetic remodeling during monolayer cell expansion reduces therapeutic potential

**DOI:** 10.1101/2021.12.14.472696

**Authors:** Adrienne K. Scott, Eduard Casas, Stephanie E. Schneider, Alison R. Swearingen, Courtney L. Van Den Elzen, Benjamin Seelbinder, Jeanne E. Barthold, Jennifer F. Kugel, Josh Lewis Stern, Kyla J. Foster, Nancy C. Emery, Justin Brumbaugh, Corey P. Neu

## Abstract

Understanding how cells remember previous mechanical environments to influence their fate, or mechanical memory, informs the design of biomaterials and therapies in medicine. Current regeneration therapies require two-dimensional (2D) cell expansion processes to achieve large cell populations critical for the repair of damaged (e.g. connective and musculoskeletal) tissues. However, the influence of mechanical memory on cell fate following expansion is unknown, and mechanisms defining how physical environments influence the therapeutic potential of cells remain poorly understood. Here, we show that the organization of histone H3 trimethylated at lysine 9 (H3K9me3) and expression of tissue-identifying genes in primary cartilage cells (chondrocytes) transferred to three-dimensional (3D) hydrogels depends on the number of previous population doublings on tissue culture plastic during 2D cell expansion. Decreased levels of H3K9me3 occupying promoters of dedifferentiation genes after the 2D culture were also retained in 3D culture. Suppression of H3K9me3 during expansion of cells isolated from a murine model similarly resulted in the loss of the chondrocyte phenotype and global remodeling of nuclear architecture. In contrast, increasing levels of H3K9me3 through inhibiting H3K9 demethylases partially rescued the chondrogenic nuclear architecture and gene expression, which has important implications for tissue repair therapies, where expansion of large numbers of phenotypically-suitable cells is required. Overall, our findings indicate mechanical memory in primary cells is encoded in the chromatin architecture, which impacts cell fate and the phenotype of expanded cells.

**SIGNIFICANCE STATEMENT:** Tissue regeneration procedures, such as cartilage defect repair (e.g. Matrix-induced Autologous Chondrocyte Implantation) often require cell expansion processes to achieve sufficient cells to transplant into an *in vivo* environment. However, the chondrocyte cell expansion on 2D stiff substrates induces epigenetic changes that persist even when the chondrocytes are transferred to a different (e.g. 3D) or *in vivo* environment. Treatments to alter epigenetic gene regulation may be a viable strategy to improve existing cartilage defect repair procedures and other tissue engineering procedures that involve cell expansion.

## INTRODUCTION

Cells constantly sense and adapt to cues from the mechanical environment, affecting cell function, differentiation, and disease states. Many studies have explored how the physical and mechanical properties of the environment affect cellular behavior ^1,2^, however, little is known about how cellular adaptations from previous physical environments are maintained, or how cells encode mechanical memory. The key to mechanical memory is to understand how physical priming in one environment will impact cellular performance and fate in another, later environment ^3–5^. In some cases, when exposure to physical priming is limited, mechanical memory can persist for a short time period (e.g. 1-3 days) and does not impact long-term cell fate ^6^. In contrast, when the exposure time to a mechanical environment increases past a critical threshold, the accumulated memory of this environment reduces cellular plasticity, resulting in cells reprogrammed to a state with a persisting phenotype ^4,7,9^ In some cases, this reprogrammed state can be maladaptive and can aggravate diseases such as fibrosis ^4,7,9^, influence cellular behavior of cancer metastasis ^10^, or limit tissue regeneration procedures ^4–6^. However, mechanisms of mechanical memory are not well understood, and investigating how to disrupt maladaptive mechanical memory for translational applications is critical.

Though mechanisms are poorly understood, recent studies focus on the organization of chromatin architecture within the nucleus ^3,5,8,11^. For example, cell stretching experiments caused chromatin condensation that persisted in a dose-dependent manner according to the time of exposure to the previous loading ^3,12^, demonstrating mechanical memory. Analyzing changes in the overall spatial organization of chromatin provides insight into changes in gene regulation, since spatial organization regulates chromatin accessibility, interactions and function ^8^. However, how the spatial organization and local gene regulation of specific epigenetic modifications retain mechanical memories has not yet been explored.

Our studies focus on the impact of mechanical memory on tissue regeneration therapies, such as cartilage defect repair procedures. Various standard tissue engineering procedures involve culturing cells on 2-dimentional (2D) tissue culture plastic (TCP) to generate sufficient cells to transplant into an *in vivo* environment. However, this expansion and associated mechanical priming alters cell fate *in vivo* ^4,13^, suggesting there is an exposure limit to *in vitro* culture expansion before the memory of this environment alters the cells’ therapeutic potential. Monolayer expansion *in vitro* is a crucial step in some cartilage defect repair procedures, such as MACI (Matrix-induced autologous chondrocyte implantation) ^14,15^. MACI involves three steps: 1) isolating a biopsy taken from a non-load bearing region of the knee, 2) expanding chondrocytes on TCP up to 16 population doublings ^16^, and 3) seeding cells onto a porcine collagen matrix for implantation into the defect ^17^. While native chondrocytes are mostly quiescent cells, the exposure to a stiff 2D substrate *in vitro* causes chondrocytes to proliferate. Although the proliferation is beneficial to generate sufficient cells for the MACI procedure, the exposure to stiff 2D substrates is known to cause chondrocytes to dedifferentiate, meaning that the hyaline (native) chondrocyte phenotype is lost and a fibroblast-like phenotype emerges ^14,18,19^. Previous work from our lab suggests that 2D culture influences the chondrocyte phenotype through changes in nuclear strain transfer ^20^. In fact, when the 3D geometry and physiological nuclear strains are restored following transfer to a 3D scaffold after dedifferentiation, it is well known that the round chondrocyte hyaline morphology returns ^21^. However, the hyaline chondrocyte phenotype is only partially rescued by changing the substrate ^14,22,23^. In this paper, we define the ability to regain the primary chondrocyte phenotype, through gene expression of chondrogenic genes, as a measure of chondrogenic potential. The decrease in chondrogenic potential due to exposure to the previous mechanical environment suggests chondrocytes retain a mechanical memory. Previous studies have not yet explored how mechanical memory could prevent proper regeneration of cartilage, motivating us to understand the extent to which chondrocytes demonstrate mechanical memory and the mechanisms of how this memory is retained.

Changes in spatial organization of chromatin or chromatin architecture could retain the persisting memory of the dedifferentiated chondrocyte state. Chromatin architecture is mediated by various epigenetic factors, such as DNA methylation or post-translational modifications of histones. The repressive histone modification, H3K9me3, is a heritable epigenetic marker that mediates cellular memory and cell fate in many cell types ^24–27^. H3K9me3 is often classified as an epigenetic barrier for cellular reprogramming ^28^, supporting that patterns of H3K9me3 have the potential to lock in cellular memory ^29–31^. Additionally, the architecture of H3K9me3 has been shown to remodel in response to changes in the cytoskeleton ^32^ and specifically in chondrocytes in response to a change in the mechanical environment ^33^. Moreover, we have shown that mechanical cues influence the reorganization of H3K9me3 and in turn direct cell fate of cardiomyocytes during development ^34^ Since H3K9me3 both restricts cellular reprogramming and is remodeled in chondrocytes and other cell types in response to a change in the biophysical environment, we hypothesized that H3K9me3 is a key regulator of dedifferentiation during expansion on 2D stiff substates that could retain a gene regulation role even in a different physical environment, thus encoding mechanical memory.

The objective of this study is to investigate the extent to which the epigenetic modification, H3K9me3, regulates the chondrocyte phenotype and retains a memory from the *in vitro* expansion process on 2D stiff substrates. This memory may limit the therapeutic potential of chondrocytes, specifically in the context of cartilage defect repair procedures. Throughout the *in vitro* culture on TCP, we showed that the chromatin architecture of regions enriched with histone modification H3K9me3 progressively changed while the chondrocyte phenotype was lost. When dedifferentiated chondrocytes were later encapsulated in a 3D environment, the gene expression profile and chromatin architecture were retained in a dose-dependent manner, suggesting a structural epigenetic memory from the previous environment. Using an inducible mouse model to suppress levels of H3K9me3, we found that decreasing levels of H3K9me3 did not prevent nuclear structural changes and did not stop the dedifferentiation process. However, by increasing the levels of H3K9me3 through inhibition of the demethylases of H3K9 (KDM4 enzymes), chondrocyte dedifferentiation was attenuated and main features of the hyaline chondrocyte chromatin structure were retained.

## RESULTS

### Histone modifications are spatially remodeled on 2D stiff substrates during expansion

To understand how the nucleus changes in response to the mechanical environment, we first analyzed the bulk nuclear morphology and intra-nuclear organization of bovine chondrocytes during expansion on a stiff 2D substrate (TCP). We confirmed the dedifferentiated phenotype reported in literature ^14,35^ by observing the cellular morphology (cell spreading and actin reorganization) and assessing the change in chondrogenic gene expression (*ACAN*, *SOX9, COL2A1* mRNA levels) compared to chondrocytes analyzed after isolation (Fig. 1A,B). As the chondrocytes dedifferentiated, we observed that the nuclear area and nuclear aspect ratio increased (Fig. 1C,D & Fig. SI 1A). By staining DNA using DAPI, we found that the chromatin architecture, spatially differed between chondrocytes after isolation (PD0) and after 16 population doublings (PD16) (Fig. 1C, middle column). Specifically, euchromatin regions (low intensity DAPI signal) were located in the center of the nucleus, while heterochromatin regions (high intensity DAPI signal) were located towards the nuclear envelope. The structural reorganization of heterochromatin and euchromatin suggested that expansion time on TCP influences mechanisms associated with regulating chromatin architecture and gene expression (e.g. histone modifications, DNA methylation).

**Figure 1.**
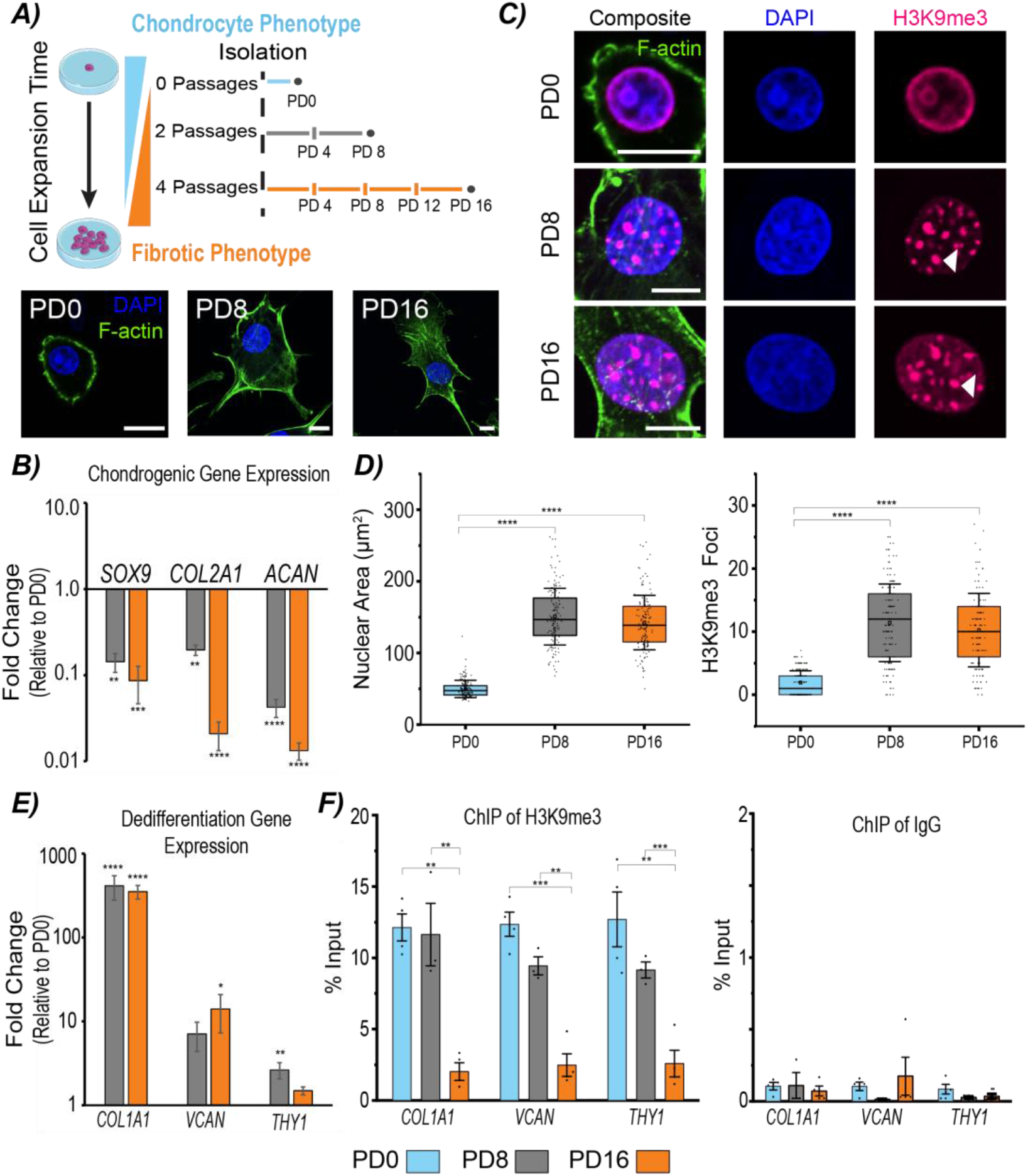
Extended culture on stiff substates leads to global remodeling of trimethylated H3K9 and a decrease in levels of H3K9me3 locally on dedifferentiation marker genes. **A)** Bovine chondrocytes were isolated and plated on tissue culture plastic (0 population doublings, PD0), passaged two times (8 populations doublings, PD8), and passaged 4 times (16 populations doublings, PD16). As populations doublings increased, chondrocytes dedifferentiated into fibroblast-like cells. PD8 and PD16 chondrocytes became more spread out (scale= 5μm). **B)** Expression of hallmark chondrogenic genes (*SOX9*, *COL2A1* and *ACAN*) decreased (±s.e.m, N=6 biological replicates). **C)** Chondrocytes expanded in a 2D stiff environment show changes in chromatin architecture reorganization (DAPI) and H3K9me3 location and distribution (scale=10μm). White indicators highlight H3K9me3 foci. **D)** With increased time on 2D stiff substrates, both nuclear area and the number of H3K9me3 foci increased (±s.d., N=6 biological replicates, n>22 nuclei/treatment/replicate). **E)** Expression of dedifferentiation markers (*COL1A1*, *VCAN*, and *THY1* mRNA levels) increased compared to PD0 cells (±s.e.m, N=6 biological replicates). **F)** Occupancy levels of H3K9me3 on the promoters of dedifferentiation genes decreased with population doublings while the background (IgG) remained low, demonstrating H3K9me3 plays a role in regulating the expression of dedifferentiation genes after PD8 (±s.d., N=3-4 biological replicates). *p<0.05, **p<0.01, ***p<0.001,****p<0.0001

As the architecture of H3K9me3 has been shown to remodel in response to changes in the mechanical environment ^33^, we studied the specific arrangement of H3K9me3 during chondrocyte dedifferentiation. We specifically focused on this repressive histone modification given that it is crucial for the regulation of cell fate ^24–27^. We first observed that H3K9me3 was localized near the nuclear envelope of the PD0 chondrocyte, but during dedifferentiation the regions of H3K9me3 formed more distinct foci distributed evenly throughout the nucleus (Fig. 1C). To quantify the overall structural shift of these epigenetic changes in the nucleus, we designed a computational method to analyze the number of H3K9me3 foci in the imaged nuclei (Fig. 1D) and found the average number of H3K9me3 foci in each nucleus increased with exposure time to the 2D stiff environment. These results indicated that the structure and pattern of H3K9me3 may play a major role in regulating chondrocyte dedifferentiation.

### H3K9 trimethylation is involved in regulating chondrocyte dedifferentiation genes

Because of the observed spatial reorganization of H3K9me3 during passaging, we explored whether this epigenetic modification mediates typical chondrocyte specific changes in gene expression during the dedifferentiation process. Using chromatin immunoprecipitation (ChIP) in conjunction with qPCR, we found that occupancy levels of H3K9me3 on promoters of chondrogenic genes (*SOX9, COL2A1, ACAN*) of PD0 cells were close to background levels, which is expected since these genes are highly expressed at PD0 (Fig. SI 2A). By PD16, the levels of H3K9me3 occupancy did not increase on the promoters of *SOX9, COL2A1*, or *ACAN*, indicating that H3K9me3 does not suppress these chondrogenic genes during dedifferentiation (Fig. SI 2A). However, we did see that the density of H3 protein increased at the promoters of these genes indicating a condensation of chromatin and potential repression of these genes through other epigenetic mechanisms (Fig. SI 2A). Additionally, we measured mRNA expression of well-established dedifferentiation genes, *COL1A1, VCAN*, and *THY1* ^14,36^, which increased expression compared to PD0 cells as the chondrocytes became more like fibroblast cells (Fig. 1E). We hypothesized that a decrease in occupancy levels of repressive marker H3K9me3 would allow for an increase in expression of these genes. We found that the occupancy levels of H3K9me3 on the promoters of chondrocyte dedifferentiation genes (*COL1A1, VCAN, THY1*) decreased after 16 population doublings compared the PD0 cells (Fig. 1F), while H3 density only decreased on the promoter of *VCAN* (Fig. SI 2B). At the same time, gene expression levels of *COL1A1* and *THY1* increased after 8 population doublings, but the occupancy levels of H3K9me3 did not significantly decrease relative to PD0 cells (Fig. 1F), suggesting that other mechanisms upregulate *COL1A1* and *THY1* of PD8 cells. Altogether, our results show the decrease in H3K9me3 occupancy levels is associated with an increase in expression of *COL1A1, VCAN*, and *THY1* after 8 population doublings on a 2D stiff substrate.

### Chromatin architecture of H3K9me3 retains an epigenetic memory from the previous mechanical environment

Next, we encapsulated PD0, PD8 and PD16 chondrocytes in 3D hydrogels and analyzed cells after 1, 5 and 10 days in 3D culture to understand if chondrocytes retained a memory from the previous mechanical environment (Fig. 2A,B). We assessed the changes in chondrogenic potential, or the ability to regain the hyaline chondrocyte phenotype (re-differentiation) ^22^, by quantifying gene expression of chondrogenic genes (Fig. 2C). In comparison to PD0-3D controls (chondrocytes encapsulated right after isolation), the chondrogenic gene expression of *SOX9* was rescued by encapsulating the cells in hydrogels for 10 days for both the PD8 encapsulated cells (PD8-3D) and PD16 encapsulated cells (PD16-3D) (Fig. 2C). In contrast, expression levels of *COL2A1* and *ACAN* were not rescued with 3D culture; expression of *COL2A1* and *ACAN* of PD16-3D cells remained significantly lower compared to PD0-3D cells even after 10 days elapsed in 3D culture (Fig. 2C). The decreased plasticity of the PD16 cells to re-differentiate suggests the changes in gene expression in 3D culture depended on the previous exposure time to a stiff 2D substrate. Thus, chondrocytes retained a mechanical memory of the previous physical environment.

**Figure 2:**
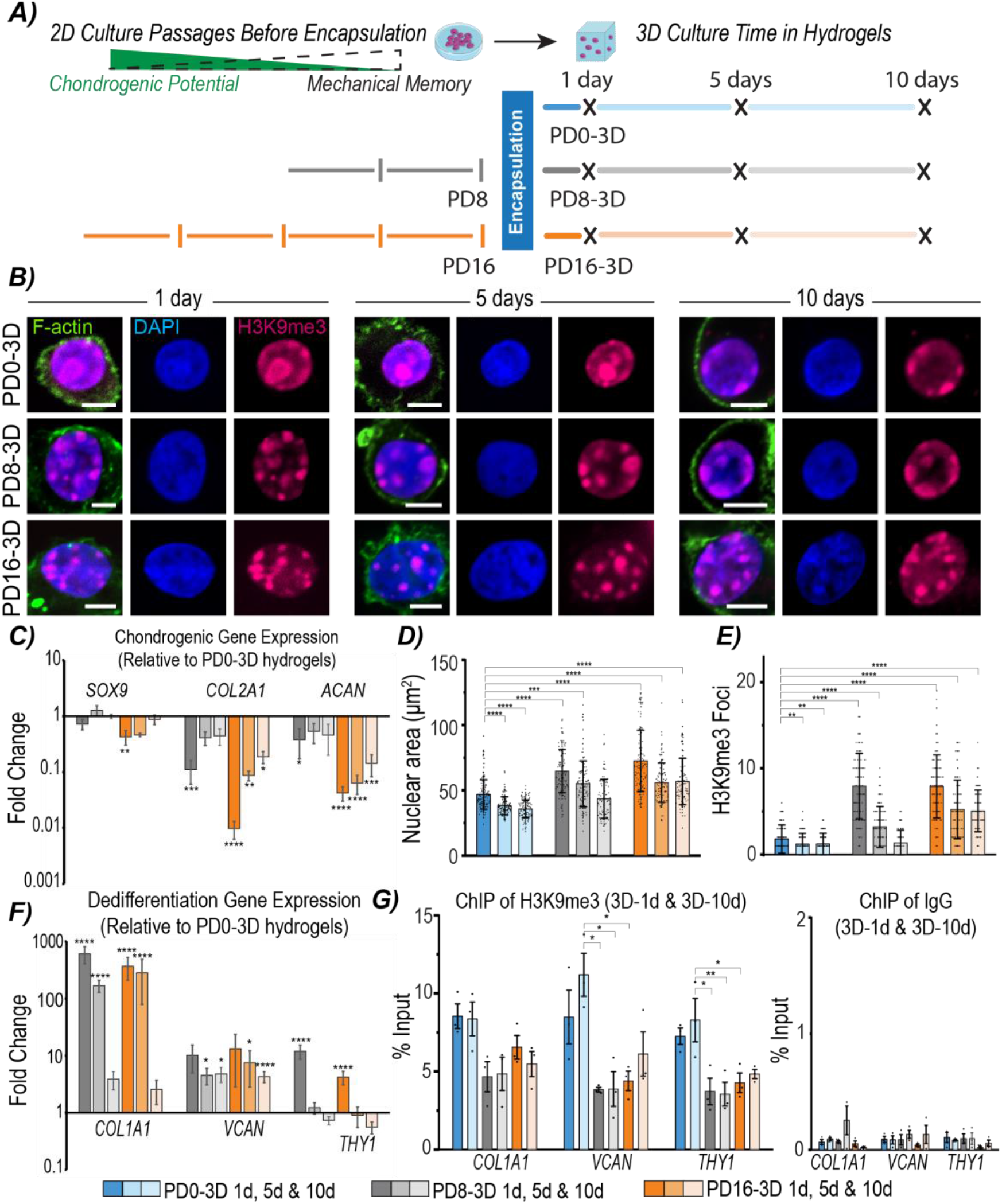
Expanded chondrocytes encapsulated in 3D hydrogels exhibited a dose-dependent mechanical memory shown by chondrogenic gene expression and H3K9me3 nuclear architecture, but not local gene regulation of dedifferentiation markers. **A)** After 0, 8, and 16 chondrocyte population doublings, cells were encapsulated into hydrogels and cultured for 1, 5 and 10 days (PD0-3D, PD8-3D, PD16-3D). **B)** Cell were stained for H3K9me3 to assess chromatin architecture adaptations in 3D culture (scale=5μm). **C)** Chondrogenic potential was assessed by changes in gene expression of chondrogenic genes of PD8-3D and PD16-3D cells compared to PD0-3D cells. Although chondrogenic gene expression of PD16-3D cells did not return to levels of PD0-3D cells after 10 days, the gene expression of PD8-3D cells did return to levels similar to PD0-3D cells after 5 or 10 days in culture, indicating a dose-dependent genetic memory from the previous culture (±s.e.m., N=6 biological replicates). **D)** Quantified nuclear area (±s.d. N=6 biological replicates, n>16 nuclei/treatment/replicate, only p-values for P0-1d comparisons are shown for clarity, Table SI 4 reports all p-values) and **E)** number of H3K9me3 foci was restored to levels of the P0-3D cells only with lower population doublings (PD8-3D cells only), demonstrating a mechanical memory from the expansion process on a stiff 2D substrate (±s.d. N=6 biological replicates, n>16 nuclei/treatment/replicate, only p-values for P0-1d comparisons are shown for clarity, Table SI 4 reports all p-values). **F)** Expression of dedifferentiation genes decreased to levels of PD0-3D cells for both PD8-3D and PD16-3D cells, however the expression of these genes did not show a dose-dependent memory (±s.e.m., N=6 biological replicates). **G)** ChIP results of PD8-3D and PD16-3D cells revealed a persistent lower occupancy level of H3K9me3 (with low IgG background) on the promoters of the dedifferentiation genes compared to PD0-3D cells, which did not change after 10 days (±s.d. N=3 biological replicates). *p<0.05, **p<0.01, ***p<0.001, ****p<0.0001

To understand how the dedifferentiated chondrocyte phenotype is retained, we next explored how changes nuclear morphology and chromatin architecture from 2D culture may be retained in 3D culture. Since we showed that the bulk nuclear properties (e.g. nuclear area and nuclear aspect ratio) changed during dedifferentiation, we investigated if the nuclear area and aspect ratio decreased in subsequent 3D culture as chondrocytes re-differentiated. We determined that after encapsulation of PD8 or PD16 cells, the nuclear aspect ratio did not return to levels of the PD0-3D cells (Fig. SI 1B). However, the nuclear area of PD8-3D and PD16-3D cells significantly decreased after 5 and 10 days in culture, potentially due to the physical constraints of the 3D environment (Fig. 2D). We confirmed that this decrease in nuclear area was not due to cell death, by imaging and analyzing the percent viable cells after encapsulation in hydrogels (Fig. SI 3A). We also found that the nuclear area of PD16-3D cells decreased over the course of 10 days (Fig. 2D, p<0.0001), but PD16-3D cells cultured for 10 days still had larger nuclei compared to PD0-3D cells encapsulated for the same culture time (Fig. 2D, p<0.0001). In contrast, the nuclear area of PD8-3D cells decreased after 10 days, approaching levels statistically similar to PD0-3D cells after 1 day in culture (Fig. 2D, p=0.2535). The increased adaptability of cells exposed to the stiff TCP for less time (PD8 cells) suggested the expanded chondrocytes retained a structural nuclear memory of the previous physical environment.

Next, we assessed the intra-nuclear architecture of H3K9me3 over time in 3D culture. The number of H3K9me3 foci of PD0-3D cells remained low over the course of 10 days (Fig. 2E). As a control, we cultured *ex vivo* cartilage plugs to confirm that the hydrogel culture was not changing the nuclear architecture of isolated chondrocytes (Fig. SI 4A). We found that the chondrocytes cultured *ex vivo* maintained similar nuclear architecture to PD0-3D cells with low nuclear area, low number of H3K9me3 foci, and low aspect ratios (Fig. SI 4B). The encapsulated PD8 and PD16 cells both had a high number of H3K9me3 foci after 24 hours. However, throughout the 10 days in culture, the amount of H3K9me3 foci of the PD8-3D cells decreased to levels statistically similar to PD0-3D cells encapsulated for 1 day (Fig. 2E, p=0.1681). At the same time, the mean number of foci observed in the PD16-3D nuclei after 10 days remained higher than the mean number of H3K9me3 foci in the PD0-3D cells encapsulated for 1 day (p<0.0001). We concluded that the organization of H3K9me3 depended on exposure time to the previous 2D stiff substrate and thus the encapsulated chondrocytes retained an intra-nuclear structural memory.

**Figure 3:**
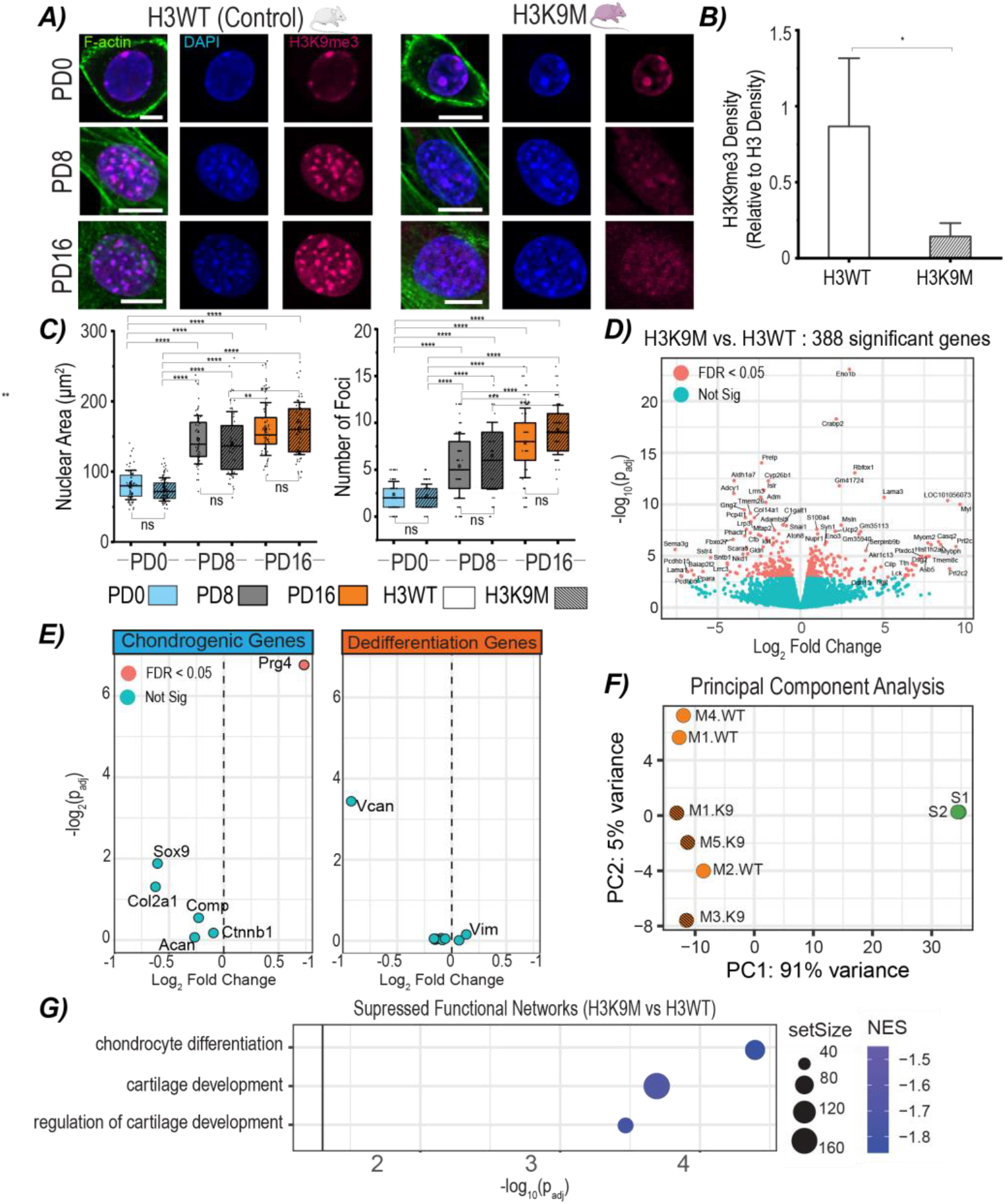
Suppression of H3K9me3 did not prevent nuclear architecture remodeling of H3K9 trimethylated chromatin and dedifferentiation of expanded chondrocytes. **A)** Chondrocytes were isolated from H3K9M mice or H3WT mice and expanded to PD16. Immunofluorescence imaging showed a decrease in H3K9me3 signal, but the nuclear architecture changes of H3K9M chondrocytes from PD0 to PD16 appeared similar to control cells. **B)** Quantified H3K9me3 density relative to total H3 density from western blots decreased in H3K9M cells compared to H3WT cells to confirm suppression of H3K9me3 (+s.d. N=4 biological replicates). *p<0.05 **C)** Nuclear area and number of H3K9me3 foci did not change significantly between H3WT and H3K9M cells of the same passage (±s.d. N=4 biological replicates, n>16 nuclei/genotype/replicate). *p<0.05, **p<0.01, ***p<0.001, ****p<0.0001, ns: p<0.05 **D)** RNA-seq analysis revealed 388 significantly differentially expressed genes. **E)** However, most chondrogenic and dedifferentiation genes were not significantly expressed. **F)** In comparison to previously published data of native chondrocytes, principal component analysis (PCA) revealed that 91% of the variance of the changes in gene expression in the system can be accounted for by the difference between the expanded and the native chondrocytes, indicating that both H3K9M and H3WT cells dedifferentiated and differ between them in a much smaller proportion. **G)** Although the lack of H3K9me3 did not prevent dedifferentiation, H3K9me3 did play a protective role in maintaining the chondrocyte phenotype since chondrogenic pathways were suppressed significantly for H3K9M cells.

**Figure 4:**
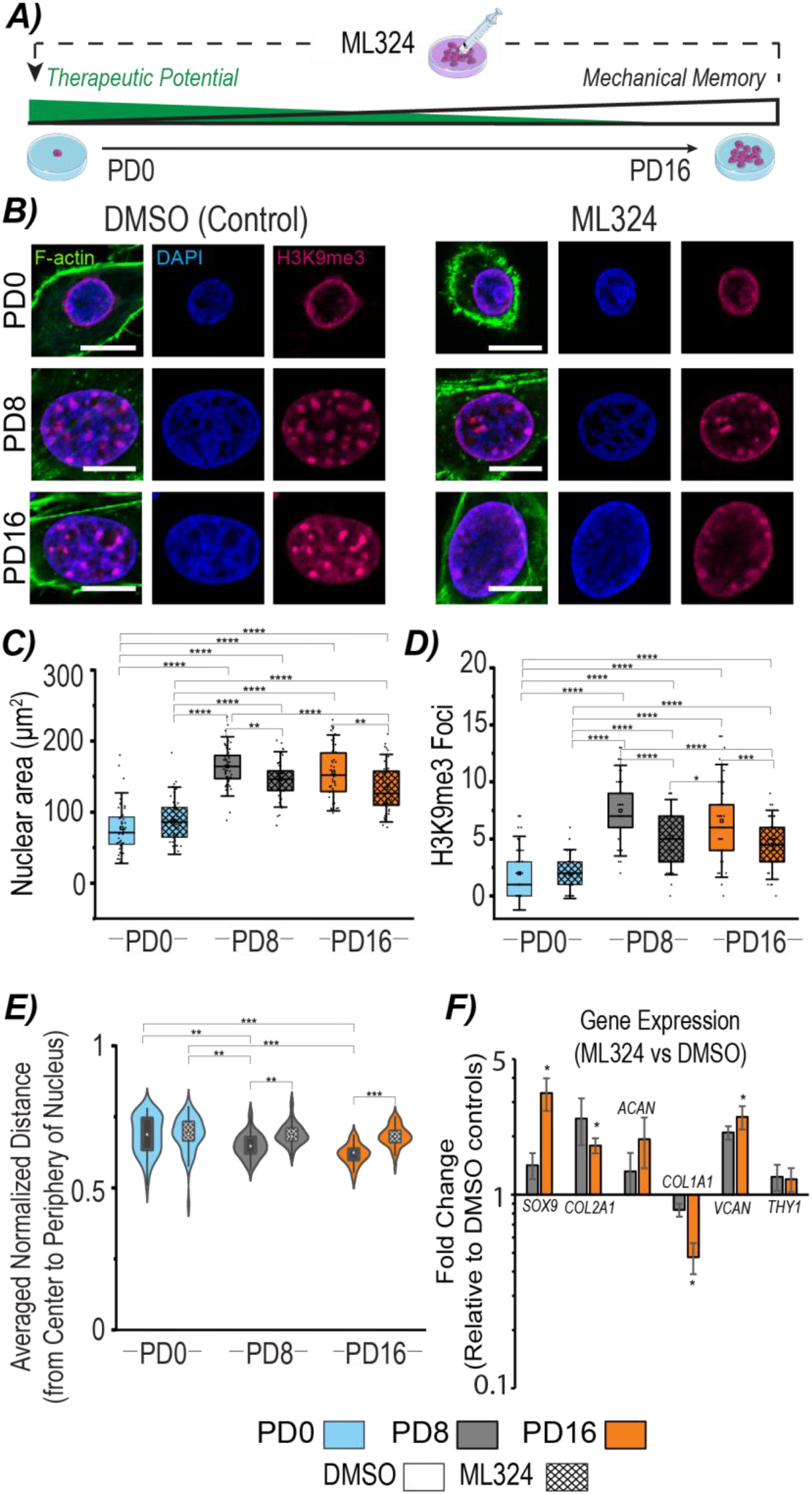
Increasing levels of H3K9me3 with treatment of a KDM4 demethylase inhibitor, ML324, partially rescued chondrogenic nuclear architecture and gene expression. **A)** During the expansion process, chondrocytes were treated with a demethylase inhibitor to increase levels of H3K9me3 and enhance the therapeutic potential of expanded chondrocytes. **B)** Chondrocytes were treated with a KDM4 demethylase inhibitor, ML324 (10 μM), or the vehicle control (DMSO) and imaged to assess nuclear architecture of H3K9me3. **C)** ML324 treatment decreased the nuclear area of PD8 and PD16 cells in comparison to PD8 and PD16 DMSO controls (± s.d., N=3 biological replicates, n>25 nuclei/treatment/replicate). **D)** ML324 treatment also decreased the number of H3K9me3 foci in comparison to DMSO controls. (±s.d. N=3 biological replicates, n>25 nuclei/treatment/replicate) **E)** The average foci location within the nucleus shifts from the periphery (1) to the center (0) of the cell to PD16, however, the ML324 treatment causes a shift in the distribution of foci location towards the periphery of the cell (box: 25^th^ to 75^th^ percentile, ±s.d. N=3 biological replicates, n>25 nuclei/treatment/replicate). **F)** With the treatment of ML324, the expression of chondrogenic genes, *SOX9* and *COL2A1*, is significantly upregulated and dedifferentiation gene, *COL1A1*, is significantly downregulated by PD16 in comparison to DMSO controls (±s.e.m. N=3 biological replicates). *p<0.05, **p<0.01, ***p<0.001, ****p<0.0001

### Dedifferentiation genes do not retain dose-dependent memory from the previous environment

Given that the epigenetic modification H3K9me3 played a role in the regulation of dedifferentiation related genes (*COL1A1*, *VCAN* and *THY1*), we hypothesized that occupancy levels of H3K9me3 would retain a memory of the dedifferentiated phenotype. When assessing the expression of dedifferentiation genes of encapsulated chondrocytes, we found that the expression levels did not show evidence of a dose-dependent memory from the previous environment. Unlike the expression of the chondrogenic genes after encapsulation (Fig. 2C), the exposure time to 2D culture prior to encapsulation did not influence the magnitude of the decrease in dedifferentiation gene expression after 10 days in the hydrogel culture (Fig. 2F). After 10 days, the expression of dedifferentiation genes of both the PD8-3D and the PD16-3D cells decreased to similar levels. However, the average change in expression of all dedifferentiation genes from 1 to 5 days in culture was greater for PD8-cells, suggesting that a memory from the previous culture may influence the initial redifferentiation rate. To further understand gene regulation of these dedifferentiation genes in the hydrogels, we performed ChIP on the 1 and 10 day samples and assessed the occupancy levels of H3K9me3 and total H3 on the promoters of the dedifferentiation genes (Fig. 2G, Fig. SI 2C). Overall, we found that the occupancy levels of H3K9me3 on the promoter of *VCAN* and *THY1* was lower for PD8-3D and PD16-3D cells in comparison to PD0-3D cells. Although gene expression of the dedifferentiation genes for PD8-3D and PD16-3D cells decreased over time, we did not find that the occupancy levels of H3K9me3 increased. Instead, we demonstrated that the levels of H3K9me3 on the promoters of *COL1A1, VCAN*, and *THY1* remained at statistically similar levels after 10 days for both PD8-3D and PD16-3D cells. However, density of total H3 protein on promoters of *COL1A1*, *VCAN*, and *THY1* of PD8-3D and PD16-3D cells increase to statistically similar levels to PD0-3D cells after 10 days (Fig. SI 2C), suggesting that other regulatory mechanisms are important for suppressing these dedifferentiation genes following encapsulation. We did conclude that the occupancy level of H3K9me3 on the promoters of *VCAN* and *THY1* depended on whether the chondrocytes were exposed to a 2D stiff environment previously since PD8-3D and PD16-3D cells had lower levels of H3K9me3 occupancy compared to PD0-3D cells (Fig. 2G). This observation suggested that epigenetic changes that occurred during dedifferentiation can be retained in 3D culture.

### Suppressing levels of H3K9 methylation did not prevent chromatin architecture remodeling or dedifferentiation

Because our findings indicated that both changes in global architecture and local occupancy levels of H3K9me3 correlate to changes in the chondrocyte phenotype, we explored how disrupting H3K9 methylation could alter cell fate and potentially alter the dedifferentiation process. We cultured chondrocytes from a mouse model with an inducible histone H3.3 lysine-to-methionine mutant (H3K9M), which acts as a global dominant negative inhibitor of H3K9 trimethylation. For controls, we used mice with an inducible expression of wild-type H3.3 (H3WT) ^37^. Unexpectedly, the overall chromatin architecture of H3K9M PD16 cells was not disrupted and remained similar to the PD16 H3WT cells (Fig. 3A,C & Fig. SI 1C). Although the fluorescent intensity of H3K9me3 (Fig. 3A) and the protein content shown by western blots (Fig. 3B & Fig. SI 5A) decreased, the number of H3K9me3 foci, nuclear area, and aspect ratio of H3K9M chondrocytes still increased with exposure time to a 2D stiff mechanical environment (Fig. 3C, Fig. SI 1C). Subsequently, we wanted to understand if suppression of H3K9 methylation led to changes in gene expression. With gene expression profiling by RNA-seq, we found significant changes in expression of 388 genes (Fig. 3D), however expression of the majority of selected chondrogenic or dedifferentiation genes (Table SI 1) of H3K9M chondrocytes did not change significantly compared to H3WT control chondrocytes (Fig. 3E). To understand if there was a difference between the dedifferentiation process of the H3K9M and H3WT chondrocytes, we compared the gene expression profiles to previously published gene expression data collected from murine articular native chondrocytes ^38^. We found that in comparison to the native articular chondrocytes, both H3K9M and H3WT cells dedifferentiated (Fig. 3F & Fig. SI 6). Principal component analysis revealed that H3K9M and H3WT cells shared similar expression profiles but clustered separately from native chondrocytes, suggesting a similar adaptation to the 2D stiff culture (Fig. 3F). Expression of chondrogenic genes (*Sox9*, *Col2a1*, *Acan*) for both H3K9M cells and H3WT cells decreased compared to native chondrocytes (Fig. SI 6A, Table SI 2,3). Dedifferentiation genes (*Col1a1*, *Col1a2*, *Thy1*, *Vim*, *Col3a1*) all increased compared to native controls (Fig. SI 6B, Table SI 2,3). *Vcan* expression of H3K9M and H3WT cells did not increase significantly compared to native controls, which is likely due to the development timing of the native chondrocyte cells, since *Vcan* is known to be more actively expressed in developing tissue compared to adult tissue ^39^. Therefore, the suppression of H3K9me3 did not prevent the chromatin architecture remodeling nor dedifferentiation of chondrocytes when exposed to stiff 2D substates, further supporting our previous findings that changes in chromatin architecture correlate to changes in the chondrocyte phenotype.

Although both H3K9M and H3WT cells dedifferentiated compared to native controls, gene expression differences suggest that H3K9me3 plays a role in regulating the chondrocyte cell fate through influencing other pathways that are not directly related to dedifferentiation. Decreased H3K9me3 in mutant histone samples altered the expression of genes that are related to cell fate commitment (Fig. SI 5B), which is expected since H3K9me3 is a critical epigenetic modification during the differentiation process ^40^. While individual dedifferentiation genes did not change significantly between H3WT and H3K9M samples, functional annotation revealed that gene expression signatures characteristic of chondrocytes were de-enriched in H3K9M samples (Fig. 3G). Assessment of the significantly changing genes in these pathways revealed several downregulated genes (e.g. *Pth1r*, *Nkx3-4*, *Grem-1*, *Snai2*, *Chadl*) that lead to an osteogenic lineage when suppressed (Fig. SI 5C) ^41–43^. Overall, our results indicate that the presence of H3K9me3 does protect a subset of specific chondrogenic functional pathways, but the suppression of H3K9me3 does not significantly influence the dedifferentiation process.

### Demethylase inhibition partially restores both chondrogenic H3K9me3 chromatin architecture and gene expression following expansion on a stiff 2D substrate

Since suppressing H3K9 methylation was not successful in altering the dedifferentiation process but decreasing the levels of H3K9me3 de-enriched some chondrogenic pathways, we hypothesized that increasing the levels of H3K9me3 by inhibiting KDM4 enzymes could disrupt dedifferentiation and increase the chondrogenic potential of expanded chondrocytes (Fig. 4A). This hypothesis is also supported by our ChIP experimental findings, since the loss of H3K9me3 occupancy levels correlated with the loss of the chondrocyte identity (Fig. 1F, 2G). When we treated cells on a 2D stiff substate with a KDM4 inhibitor, ML324 ^44^, during the expansion process, we found that chromatin architecture (H3K9me3 foci, nuclear area) changed significantly in comparison to the vehicle control (DMSO) cells (Fig. 4B,C,D), yet the nuclear aspect ratio did not change significantly compared to DMSO controls (Fig. SI 1D). We found that not only did the number of foci change, but also the location of the foci in the PD16 treated cells. Most foci in the treated cells appeared to be concentrated around the periphery of the nucleus rather than evenly throughout the nucleus like the DMSO controls (Fig. 4B). To quantify the shift in location of the foci we designed a computational method to determine the average normalized distance of all foci, from the center of the nucleus (value of 1 means the foci is found toward the nuclear envelope, while a value of 0 means the foci is at the center of the nucleus). We found that the average distance of foci from the periphery to the center is lower for PD8 and PD16 DMSO control cells (Fig. 4E). However, with the ML324 treatment, the average foci location from the center is higher and statistically similar to the PD0 cells (Fig. 4E). Additionally, ML324 treatment led to a significant increase in expression of *SOX9* and *COL2A1* and a decrease in expression of *COL1A1*, but did not decrease the expression of dedifferentiation gene *VCAN* (Fig. 4F). In contrast to decreasing levels of H3K9me3, we found that increasing levels of H3K9me3, by inhibiting KDM4 enzymes, partially rescued the characteristic native chondrocyte chromatin architecture and led to higher expression of important chondrogenic genes. Taken together, our results suggest that increasing levels of H3K9me3 may protect the chondrogenic phenotype by maintaining the nuclear architecture even when cultured on stiff 2D substrates.

## DISCUSSION

Our work demonstrates how cellular adaptations to the mechanical environment can be encoded in altered gene regulation through epigenetic modifications. As previously demonstrated in the literature, we observed that longer exposure times to stiff mechanical environments led to decreased cellular plasticity when transferred to new mechanical environments. In our experiments, the plasticity, or ability for cells to change phenotypes, was reflected by alterations in the chromatin architecture of the epigenetic modification H3K9me3. In response to a 2D stiff mechanical environment, the chromatin architecture of chondrocyte cells remodeled. When the chondrocytes were transferred to 3D hydrogels, the nuclear architecture persisted in a dose-dependent manner according to exposure time to 2D stiff environment, demonstrating a structural nuclear memory of H3K9 trimethylated chromatin. Notably, the chromatin architecture of cells exposed to a 2D stiff substrate for 16 population doublings, persisted for 10 days. We also found that the change in gene expression over a specified time in 3D culture was greater for PD8-3D than PD16-3D cells, supporting the idea that PD8-3D cells are more able to adapt quickly. Therefore, transfer to a 3D environment only rescued the chondrogenic potential of cells that had been exposed to a stiff 2D substrate for less time (as in the case of PD8 cells). To partially prevent the chromatin remodeling of cells that have been exposed to a 2D substrate for longer time periods (PD16 cells), an epigenetic treatment was needed, highlighting a potential strategy for procedures that require cell expansion processes.

The decreased plasticity due to an exposure to a stiff 2D mechanical environment suggests there is a limit to 2D expansion time before cells can no longer adapt to new environments. Specifically, for cartilage defect repair, these data suggest that after expanding chondrocytes for 16 population doublings, the chondrogenic potential, measured by changes in gene expression of chondrogenic genes, will decrease. However, why there is a dose-dependent effect from the external environment is still unknown. Perhaps, over time, changes in more stable epigenetic modifications occur until cells reach a more permanent state, similar to a ball becoming more stable while rolling down into a valley of Waddington’s landscape ^45^. For example, we found as chondrocytes dedifferentiated, the occupancy levels of H3K9me3 decreased by PD16 on dedifferentiation genes *COL1A1*, *VCAN*, and *THY1*. However, the levels of H3K9me3 was not lost at PD8, while the expression of these genes already increased. It’s possible that the loss of H3K9me3 is a more stable epigenetic change, suggesting the PD16 cells are less plastic. For the PD8 cells, other gene regulation could have been at play to increase expression and these mechanisms of gene regulation could be less stable. Understanding how cellular plasticity is controlled by the mechanical environment through changes in epigenetic factors could provide information to reverse the effects of cell expansion or optimize cellular programming for tissue regeneration procedures.

One notable change in the nucleus of the dedifferentiated chondrocyte was the increased number of H3K9me3 foci. From a biophysics perspective, the formation of these heterochromatin foci could be driven by liquid-liquid phase separation, in which interactions between multivalent macromolecules can cause the formation of separated compartments ^46^. Functionally, these foci regions may represent chromocenters, composed of clusters of pericentromeres, and characterized by methylated DNA and the occupation of H3K9me2/me3 that are bound to HP1 ^47^, but we did not confirm this experimentally. Since centromeres are responsible for correct chromosome segregation during cell division ^47^, it would make sense that these chromocenters may be more evenly distributed and accessible for chromatin organization as the number of population doublings increases because chondrocytes are known to divide much more quickly on 2D TCP environments when compared to 3D environments or *in vivo* ^22^. Perhaps the increased cell division requires reorganization of chromatin architecture to distribute chromocenters throughout the cell for efficient segregation of chromosomes.

Along with the increase in number of H3K9me3 foci with population doublings, we also observed a decrease in occupation of H3K9me3 towards the nuclear envelope, which could also be responsible for phenotypic changes seen during dedifferentiation. The localization of H3K9me3 in the Lamina-associated domains (LADs) is characteristic of fully differentiated cell types ^48^. Throughout differentiation, facultative heterochromatin dynamically associates to the nuclear envelope in the LADs, generally repressing genes that are not necessary to certain cell types ^48^. In addition to gene repression, LADs play a critical role in organizing the genome throughout the nucleus ^8^. In fact, when these domains are disrupted, variable reactivation of genes can occur often leading to a maladaptive or diseased state. For example, evidence suggests that disruption of heterochromatin near the nuclear envelope is a characteristic of many tumor cells ^49,50^. Additionally, our lab has shown that a loss of heterochromatin accumulated toward the periphery of the nucleus is a transition that occurs in cardiac hypertrophy ^34^. Therefore, the disruption of the peripheral enrichment of H3K9me3 modified chromatin could be causing aberrant gene expression of the dedifferentiated chondrocyte and a deviation from the hyaline native chondrocyte phenotype. However, further FISH or ChIP experiments would be necessary to demonstrate whether there is a change in location of dedifferentiation genes from the periphery to an active area of transcription.

The chromatin remodeling and the disruption of the LADs in response to the 2D stiff environment could be partially due to the changes in cytoskeleton structure during dedifferentiation. Mechanical strain can cause cytoskeleton remodeling, increased accumulation of nuclear envelope proteins such as emerin, and global chromatin rearrangements including decreased localization of H3K9me2/me3 in the LADs ^32^. Because the LADs are connected to the cytoskeleton through the LINC complex, the reorganization of LADs in response to changes in the cytoskeleton is plausible. Additionally, previous work disrupting the cytoskeleton of chondrocytes has been beneficial to attenuate the dedifferentiated state ^51,52^. We postulate that the shift in chromatin architecture could partially be due to the drastic changes in the cytoskeleton that are known disrupt the organization of H3K9me3. However, when chondrocytes are encapsulated the cellular morphology changes and the actin cytoskeleton remodels again. Our experiments show that epigenetic changes persist even with the change in cytoskeleton structure, suggesting other interventions are also necessary to control the chondrocyte phenotype when encapsulated in 3D.

For this reason, we explored the effect of an epigenetic treatment on chondrocytes during the dedifferentiation process. Epigenetic treatments are increasingly being approved by the FDA, supporting the feasibility of using epigenetic treatments clinically ^53–56^. Additionally, many epigenetic treatments have been shown to be useful to direct the chondrocyte phenotype ^15,57^. We found that the demethylase inhibitor, ML324, altered the chromatin architecture to appear more similar to that of native hyaline chondrocytes, despite the persistent biophysical cues from the *in vitro* expansion process, suggesting epigenetic treatments may override biophysical cues. However, the chondrocyte phenotype was not completely restored to the original native state. Future work should be done to assess the advantage of ML324 and other epigenetic treatments in combination with a return to 3D culture to increase the therapeutic potential of cultured chondrocytes. Elucidating the epigenetic mechanisms of mechanical memory would identify pharmacological targets to reprogram cells to respond in an optimal way to the surrounding environmental and chemical cues.

Taken as a whole, our results reveal the influence of the mechanical environment on cellular plasticity and the role of chromatin remodeling during dedifferentiation and in response to physical cues. An enigma of chromatin regulation is how epigenetic changes provide both stability and cellular plasticity to internal and external signals. Our work addresses this question, showing that exposure time to a stimulus plays a major role in determining stability and plasticity of cellular responses. Overall, a deeper understanding of how the physical environment influences chromatin remodeling and gene regulation could both improve strategies for tissue regeneration and provide critical information about diseases and development.

## MATERIALS AND METHODS

We expanded primary bovine chondrocytes on TCP for various passages and then encapsulated the cells in 3D hydrogels to mimic transplantation of chondrocyte cells into a 3D environment for cartilage defect repair. While growing the chondrocytes on TCP, we analyzed the cultured cells before the first passage (0 population doublings: PD0), after passage two (8 population doublings: PD8), and after passage four (16 population doublings: PD16). Additionally, PD0, PD8, and PD16 cells were encapsulated in PEGDA-HA hydrogels to study redifferentiation in 3D culture and how cells might retain a memory from the previous environment. We analyzed encapsulated samples after 24 hours, 5 days, and 10 days in 3D culture to investigate short-term and long-term cellular changes (Fig. 2A). Cellular and nuclear morphology, gene expression, and gene regulation were assessed for all samples. To understand if we could prevent the epigenetic changes, we both decreased and increased levels of H3K9me3. First, we suppressed levels of H3K9me3 by expanding cells isolated from a mouse model with an inducible histone H3.3 lysine-to-methionine mutant (H3K9M), which acts as a global dominant negative inhibitor of H3K9 methylation. Gene expression was evaluated through analysis of RNA-seq data. Next, we increased levels of H3K9me3 by treating expanded chondrocytes with ML324. Samples were analyzed using immunofluorescence and RT-qPCR. For complete details of materials and methods used, please refer to SI.

## ACKNOWLEDGEMENTS

We are grateful for the support from Dr. Thomas Cech and would like to thank the Cech lab for sharing their protocol and resources to learn the chromatin immunoprecipitation process. Additionally, we would like to thank the Genomics Shared Resource Facility at the UC Anschutz Medical Campus for library preparation and RNA sequencing of our samples. This work was supported in part by grants to C.P.N. NIH R01 AR063712 and NSF CMMI CAREER 1349735. J.B. is grateful for support from the NIH R35 GM142884 and Boettcher Foundation’s Webb-Waring Biomedical Research Awards program. S.E.S was funded through NIH T32 GM-065103. J.L.S. was supported by a postdoctoral fellowship 127621-PF-16-099-01-DMC from the American Cancer Society.

## AUTHOR CONTRIBUTIONS

Conceptualization, A.K.S. and C.P.N.; Methodology, A.K.S, E.C., S.E.S, B.S., J.F.K, J.L.S, J.B., and C.P.N.; Formal Analysis, A.K.S. and E.C.; Investigation, A.K.S, E.C., S.E.S, A.R.S, J.E.B, J.L.S, K.J.F; Statistical Analysis: C.L.V. A.K.S. and N.C.E. Writing – Original Draft, A.K.S.; Writing – Review & Editing, All authors; Funding Acquisition, C.P.N.

## Supplemental Information

### SI Materials and Methods

#### 2D Cell Culture of Bovine Chondrocytes

We harvested chondrocytes from cartilage extracted from juvenile bovine stifle (2-week-old knee) joints within 12 hours of slaughter. Using aseptic techniques ^58,59^, we harvested the cartilage of the femoral medial condyles ensuring the cells from each animal were kept separate. We rinsed obtained tissue 3 times with sterile PBS (Hyclone, Cat. No. SH30028.03) and digested the tissue with 0.2% collagenase-P (Roche Pharmaceuticals, 11213873001) for 5-6 hours at 37°C with agitation (230 rpm). After the digestion, we quenched the digested tissue with media and filtered with a 70 μm filter (Fisher, Cat. No. 22363548) to remove extracellular material and isolate chondrocytes. Chondrocyte media consisted of chemically defined DMEM/F12 (Gibco, Cat. No. 11330-032) supplemented with 10% Fetal Bovine Serum (Gibco, Cat. No. 26140-079), 0.1% bovine serum albumin (Sigma, Cat. No. A9418-100g), 1X Pen/Strep (Gibco, Cat. No. 15140-122) and 50 μg/mL ascorbate-2-phosphate (Sigma, Cat. No. 49752-10G) ^59^. We seeded chondrocytes on tissue culture plastic (TCP; Corning, Cat. No. 430167 or Ibidi Cat. No. 8084) at a density of 4×10^4^ cells/cm^2^ and expanded the cells for up to 16 population doublings (PD16). Cells were passaged around 80% confluency. All samples were incubated at 37°C and 5% CO_2_ in chondrocyte media.

#### 3D Cell Culture of Bovine Chondrocytes

We encapsulated chondrocyte cells after isolation (0 population doublings, PD0), passage 2 (8 population doublings, PD8) and passage 4 (16 population doublings, PD16) in 3D hyaluronan-poly(ethylene) glycol diacrylate (HA-PEGDA) hydrogels and cultured for 1, 5 and 10 days. To make HA-PEDGDA hydrogels, we lyophilized 23-25% thiolated hyaluronan (HA, Lifecore Biomedical, Cat. No. HA100k-5), dissolved the HA in sterile PBS, and combined the HA with a PEGDA (Alfa Aesar, Cat. No. 46497) solution ^60^. We used a final concentration of 20 mg/1mL of HA and 8.6 mg/mL PEGDA, resulting in a ratio of 1:0.8 thiol groups to PEGDA. In the HA/PEGDA gel, we encapsulated cells at a concentration 20×10^6^ cells/mL in 100uL gels. Next, we cultured the encapsulated cells at 37°C for 30 minutes to facilitate the crosslinking between the diacrylate groups on the PEGDA molecule and the thiol groups on the thiolated HA molecule, and then further maintained at 37°C and 5% CO_2_ in chondrocyte media. To name each sample, we abbreviated encapsulated cells by the number of population doublings undergone before encapsulation and denoted these samples with “3D” because they were cultured in a 3-dimentional culture. For example, cells encapsulated in PEGDA-HA hydrogels after 8 population doublings on TCP are abbreviated as PD8-3D cells. All experimental time points include: PD0, PD8, PD16, PD0-3D 1day, PD0-3D 5days, PD0-3D 10days, PD8-3D 1day, PD8-3D 5days, PD8-3D 10days, PD16-3D 1day, PD16-3D 5days, PD16-3D 10days.

#### 2D Cell Culture of H3K9M and H3WT Murine Chondrocytes

For H3K9M and H3WT (control) studies, male mice carrying the transgene ^37^ were bred to C57BL/6 (Jackson Laboratory, Cat. No. 000664) females mice. Chondrocytes from embryonic (E18.5) H3K9M and H3WT mice were harvested and cultured with chondrocyte media and 2μg/mL doxycycline (Sigma, Cat. No. D3072-1ml) for 0, 8, 16 population doublings (PD) on TCP (Corning, Cat. No. 430165 or Ibidi Cat. No. 80841). To harvest embryonic chondrocytes, hind limb articular cartilage was isolated and washed 3× with PBS, transferred to 3mg/mL collagenase-P solution in DMEM F12 (with 1X Pen/Strep), digested for 12hrs at 37C, and filtered through a 70 μm mesh strainer, and plated at a density of 4×10^4^ cells/cm^2 61^.

#### Immunofluorescence Staining & Imaging

We fixed cells cultured on TCP in 4% paraformaldehyde (PFA, Electron Microscopy Sciences, Cat. No. 15714-S) for 10 minutes, permeabilized in 1% Triton-X100 (Sigma, Cat. No. 78787-100mL) in PBS for 10 minutes. Before incubating with the primary antibodies, we blocked by incubating the cells with 10% Natural Goat Serum (NGS, Invitrogen, Cat. No. 10000C), 1% bovine serum albumin (BSA) in 0.1% PBT (0.1% Tween-20, BioRad, Cat. No. 170-6531, in PBS) for 60 minutes at room temperature (RT). We performed the primary antibody incubation in 0.1% PBT containing 1% BSA at 4°C overnight (12 hours) and the H3K9me3 primary antibody (abcam, ab8898, 1:600) with agitation. Following the primary incubation, we performed the secondary incubation in 0.1% PBT containing 1% BSA, with Alexa Fluor 633 goat anti-rabbit IgG (Life Technologies, Cat. No. A21070) at a dilution of 1:500 for 45min (RT) with agitation. To visualize both actin and DNA, we counterstained with 488 Phalloidin (Invitrogen, Cat. No. A12379, 1:80, 20 minute incubation in PBS) and DAPI (Invitrogen, Cat. No. D1306, 1:1000, 10 minute incubation in PBS) respectively.

To stain cells encapsulated in 3D HA-PEGDA hydrogels, we used a similar protocol. However, we fixed the cells in 4% PFA for 30 minutes, permeabilized in 1% Triton-X100 in PBS for 15 minutes, and blocked with 5% NGS, 1% BSA in 0.1% PBT for 60 minutes. The primary antibody incubation was performed in 0.1% PBT containing 1% BSA at 4°C for 16 hours with agitation. We used the same primary and secondary antibodies listed previously. Secondary incubation was also performed in 0.1% PBT containing 1% BSA, at a dilution of 1:200 for 2 hours (RT) with agitation. Lastly, we counterstained both actin and DNA with 488 Phalloidin (1:80, 40 minute incubation in PBS) and DAPI (1:500, 15 minute incubation in PBS) respectively.

Using an inverted Nikon A1R Confocal microscope with a 60x oil immersion objective, we imaged both encapsulated cells and cells grown on TCP from all timepoints, treatments and genotypes (Fig. 1 & 2). Specifically, we imaged the nucleus/DNA (DAPI; 405nm), actin (phalloidin; 488nm), and H3K9me3 (640nm).

#### Image-based Structural Nuclear Analysis

Using a custom MATLAB (R2020a) code, we analyzed imaged nuclei from each experimental group (n<15 nuclei/experimental group/animal). For each nucleus, the nucleus was segmented using a reference (DAPI, 405 channel) signal, and intensity values were normalized according to previously established methods ^34^. The nuclear area and aspect ratio were calculated using the MATLAB image processing toolbox. H3K9me3 foci were detected by finding the local maxima in each segmented nucleus in the H3K9me3 signal (640 channel) using the MATLAB script FastPeakFind (v. 1.13.0.0) created by Adi Natan in 2021 ^12^. Detected peaks were counted as H3K9me3 foci. To track the change in location of the foci for the cell treated with ML324 or DMSO, the distance to the center of the nucleus (*a*) and distance from the nuclear periphery (*b*) was calculated for each detected H3K9me3 foci. The ratio of *a/(a+b)* was calculated to determine the relative location of each foci relative to the center of the nucleus. MATLAB code is available upon request from the corresponding author.

#### Gene Expression assessed with RT-qPCR

We quantified expression of previously established genes that change during chondrocyte dedifferentiation (*COL2A1*, *COL1A1*, *ACAN*, *SOX9*, *VCAN*, *THY1*). Total RNA was isolated from both 3D and 2D cultures to assess relative gene expression using Real-Time quantitative PCR (RT-qPCR). To isolate RNA from 3D cultures, we homogenized the HA-PEGDA matrices using a Tissue Ruptor for 30 seconds in QIAzol lysis Reagant (Qiagen, Cat. No. 79306) ^62^. Lysed cells from 2D cultures on TCP were collected in QIAzol. After lysis, we extracted the total RNA with the E.Z.N.A. Total RNA kit (Omega Tek, Cat. No. R6834-01) or the Direct-zol RNA MiniPrep (Zymo Research, Cat. No. R2050). To reverse transcribe the isolated RNA to cDNA, we used iScript Reverse Transcription Supermix (BioRad, Cat. No. 1708841). RT-qPCR was performed using Sso AdvancedTM Universal SYBR^®^ Green Supermix (BioRad, Cat. No. 1725271) in a CFX96 Touch adthermocycler (BioRad). We designed all primers (sequences listed in Table S5) using NCBI primer blast, ensuring that all primers spanned an exon-exon junction, were specific for all known splice variants, and efficiency of all primers was calculated to be above 90%. Integrated DNA Technologies (IDT) performed the synthesis of primers. We normalized all data to the average of reference genes, HPRT1 and GAPDH. To evaluate the relative change in gene expression, we used the ΔΔCt method.

#### RNA-seq Gene Expression

H3K9M and H3WT chondrocytes were cultured in the presence of doxycycline (2 μg/mL) for 16 population doublings, lyzed with Qiazol, and RNA was extracted using Direct-zol RNA MiniPrep (Zymo Research, Cat. No. R2050). Poly A selected RNA was sequenced at the Genomics Shared Resource Facility at UC Anschutz Medical Campus. We mapped reads using hisat2 v2.1.0 ^63^ and custom parameters to the UCSC mouse genome release mm10. We summarized reads to genes annotated by UCSC using featureCounts v1.6.2 ^64^. We further assessed count data in R 3.6 using DESeq2 ^65^ to analyze differentially expressed genes and used clusterProfiler ^66^ for the gene set enrichment analysis of Gene Ontology pathways. We compared our data to publicly available RNA-seq data of embryonic proliferating chondrocytes ^38^ using in-house scripts based on DeepTools ^67^ and the bamTools and bedTools suite. Our RNAseq data is available on the GEO database (GSE190339).

#### Chromatin Immunoprecipitation (ChIP)

For adherent chondrocyte cells cultured on 2D, we crosslinked 2×10^6^-4×10^6^ cells in 1% PFA in PBS for 10 minutes, quenched in 150 mM glycine (Fisher, Cat. No. BP381-500), and scraped to harvest the cells. To harvest cells from the HA-PEGDA hydrogels, we followed a protocol for harvesting tissues for ChIP ^68^. Briefly, we homogenized the hydrogels using a disposable plastic pestle in ice cold PBS. Then, we crosslinked the cells with 1% methanol-free formaldehyde for 15 minutes with agitation and quenched in 150mM glycine. Next, we harvested the hydrogels by centrifugation at 2000g for 10mins.

We lysed both 2D and 3D cultured cells in lysis buffer, 50 mM Tris-Cl (Sigma, 93363-50G), 10mM EDTA (Fisher, Cat. No. BP120-1), 0.5% SDS (Sigma, Cat. No. L3771-100g), 1× Protease Inhibitor (Sigma, Cat. No. SRE0055-1BO). Then, we sheared the chromatin by sonicating the samples in a Bioruptor UCD-200 (Diagenode) or a M220 Focused Ultrasonicator (Covaris) so that fragment sizes of chromatin were between 200 bp to 500 bp. After shearing, we added 20 μg of chromatin to IP buffer (16.7 mM Tris-Cl, 1.2 mM EDTA, 167 mM NaCl (Fisher, Cat. No. S271-500), 1% Triton X-100). We performed Chromatin Immunoprecipitation (ChIP) on both cells cultured in 3D and on 2D ^69,70^ with 2 μg of ChIP validated antibody for the protein of interest, H3K9me3 (abcam, ab8898), as well as negative control non-specific IgG (Millipore, Cat. No. 12370) and a positive control H3 (abcam, Cat. No. ab1791) antibodies. Antibody chromatin complexes were incubated overnight with rotation. Protein A/G magnetic beads (Pierce, Cat. No. 88803) were then added for 3 hours at 4°C with rotation. We washed the beads with a series of buffers: low Salt Buffer (20 mM Tris-Cl, 2mM EDTA, 150 mM NaCl, 0.1 % SDS, 1% Triton X-100), High Salt Buffer (20 mM Tris-Cl, 2mM EDTA, 500 mM NaCl, 0.1% SDS, 1% Triton X-100), LiCl Buffer (10mM Tris-Cl, 1mM EDTA, 250 mM LiCl; Sigma, Cat. No. L9650-100G), 1% Sodium Deoxycholate (Sigma, Cat. No. D2510-100G), 1% IGEPAL (Sigma, Cat. No. 18896-50ML)). We eluted the chromatin in elution buffer (1% SDS, 100mM NaHCO_3_; Fisher Cat. No. S233-500), and added 5 uL of 5 M NaCl and to reverse the crosslinking during an overnight incubation at 65°C. Protein and RNA was digested with Proteinase K (Ambion Life Technology, Cat. No. AM2542) and RNAse A (Ambion Life Technology, Cat. No. AM2272) before the DNA was purified with either DNA clean and concentrator kit (Zymo, Cat. No. D5201) or phenol chloroform (VWR, Cat. No. 97064-692) extraction and ethanol precipitation. We analyzed the purified DNA using qPCR with primers designed to amplify the promoter regions of the following genes, *COL1A1*, *THY1*, *VCAN*. We designed the DNA primers (Table SI 6) using NCBI primer blast and ordered the primers from IDT. To determine the relative enrichment of each protein of interest for the specified promoter regions, we calculated the percent input, or the signal of an IP sample relative to the qPCR signal from the input samples (samples without IP).

#### Western Blot

Following 16 population doublings, H3K9M and H3WT chondrocytes were prepared for western blot analysis by nuclear isolation following a previously reported protocol ^34^. Briefly, cells were resuspended in cold nuclear isolation buffer (50 mM Tris-HCl pH 8, 15 mM NaCl, 60 mM KCl, 5 mM MgCl2, 1 mM CaCl2, 250 mM sucrose, 1 mM dithiothreitol, 0.6% IGEPAL (Sigma-Aldrich)) supplemented with complete protease inhibitors (Sigma-Aldrich) and incubated for 5 minutes on ice. Isolated nuclei were then centrifuged (960 × g for 5 min), washed in nuclear isolation buffer and lysed in RIPA buffer (50 mM Tris-HCL (pH 8), 150 mM NaCl, 0.1% SDS, 0.5% sodium deoxycholate, 1% Triton X-100 and 1 mM EDTA (all from Sigma-Aldrich) supplemented with complete protease inhibitors (Sigma-Aldrich) and 0.01 U μl–1 benzonase (Novagen). We sonicated the resulting lysates 10 times for 30 seconds with a 30 second pause between pulses using a Bioruptor Pico sonicator (Diagenode). The lysates were then cleared to remove cell debris through centrifuging and collecting the supernatant which was boiled together with Laemmli sample buffer (100 mg ml–1 SDS, 250 mM Tris pH 6.8, 1 mg ml–1 bromophenol blue and 50% glycerol (all from Sigma-Aldrich) and loaded into 4–20% mini-Protean TGX precast protein gels (BioRad). Protein was transferred to PVDF membranes (Bio-Rad) and blocked for 1 h in 5% powdered milk in Tris-buffered saline and Tween-20. The following primary antibodies were used: H3K9me3 (Abcam, 8898; 1:1,000), H3 (Abcam, 1791; 1:10,000 dilution). Goat, anti-rabbit-HRP-conjugated (Invitrogen, PI31460; 1:2,000 dilution) was used as the secondary antibody. Immobilon western chemiluminescent HRP substrates (Millipore) were used to detect proteins.

#### ML324 Treatment

Twenty-four hours after we seeded the isolated chondrocytes (PD0) and expanded (PD8 and PD16) chondrocytes, we treated the cells with chondrocyte medium supplemented with 10 μM of ML324 ^71^ (MedChemExpress, Cat. No. HY-12725) or a carrier control DMSO (Sigma, Cat. No. 276855-100mL). All inhibitors were dissolved in DMSO and the final concentration of DMSO in medium was maintained below 0.1%, which has been shown not to change basic metabolic activity and cell proliferation ^52^ and we confirmed that the DMSO did not significantly affect cell viability (Fig. SI 3B). Control culture medium contained 0.1% DMSO. We collected cells for imaging and gene expression analysis on PD0 (24hrs after isolation, PD8 and PD16.

#### Statistical Analysis

All statistical testing was performed using R (version 4.0.3) and Rstudio (version 1.3.1093) software. All datasets (number of foci, nuclear area, nuclear aspect ratio, chromatin immunoprecipitation results, gene expression results, foci location) were analyzed using a linear mixed effect model (nlme package, version 3.1-149) with a type II or III sum of squares ANOVA using the containment method for estimating denominator degrees of freedom (car package, version 3.0-10). Type III ANOVA was used when the model contained an interaction term (Fig. 4 C, D, E, F) and a type II ANOVA was used for all other linear models. Each model contained Passage and/or Treatment as fixed effects and we allowed the model intercept to vary with the experimental animal to control for difference among animals. Normality of the residuals was evaluated using the Shapiro-Wilk test for normality and, if necessary, the response was transformed to meet the assumptions of the model. Specifically, nuclear area measurements (Fig. 1D, 2D, 3C, 4C & Fig. SI 4B) were transformed using a square root function (*measurement*^(1/2)^), nuclear aspect ratio measurements (Fig. SI 1,4B) were transformed using a natural log function (ln(*measurement*)) and the foci location data was transformed using a arcsine-square root transformation (arcsin(*measurement*^(1/2)^)). All Shapiro-Wilk test statistics (*W*) values for the final models were greater than 0.9, meeting the assumption of normality of residuals. Additionally, homogeneity of the residual variance was evaluated for each data set by plotting the residuals against the fitted values of the linear model. In some models (number of foci: Fig. 1D, 2E, 3C, 4D, nuclear area: Fig. 1D, 2D, 3C, 4C & Fig. SI 4B, nuclear aspect ratio: Fig. SI 1,4B) the variance was dependent on the passage treatment level, so a heterogenous variances model was used to allow for differences in variance among the groups. The use of the heterogenous variance model was also justified when the likelihood ratio test showed a significant difference (*P*>0.05) between the heterogenous variance model and the linear mixed model with constant variance. We then used the emmeans function (package emmeans, version 1.5.3) to calculate the estimated marginal means and test whether each treatment levels was statistically significantly different from each other level, accounting for the differences in variance. The containment method was again used for estimating denominator degrees of freedom. P values were adjusted for multiple comparisons using the Tukey method. A P-value less than 0.05 was considered significant, and significance was denoted as \**P*<0.05, \*\**P*<0.01, and \*\*\**P*<0.001, \*\*\*\**P*<0.0001. Note the linear model of data with low numbers of samples may have too low of power to determine accurate *P*-values when the effect size is small (Fig. 1F, comparison between PD0 and PD8). However, when the effect size is large enough (Fig. 1F, comparison between PD0 and PD16), the *P*-values are considered accurate.

**Table SI 1.** List of chondrogenic and dedifferentiation genes assessed within Figure 3E.

| Phenotype | Figure 3e Genes |
| --- | --- |
| Chondrogenic | *Sox9* |
| Chondrogenic | *Col2a1* |
| Chondrogenic | *Acan* |
| Chondrogenic | *Ctnnb1* |
| Chondrogenic | *Prg4* |
| Dedifferentiation | *Col1a1* |
| Dedifferentiation | *Thy1* |
| Dedifferentiation | *Vcan* |
| Dedifferentiation | *Col1a1* |
| Dedifferentiation | *Col3a1* |
| Dedifferentiation | *Col10a1* |
| Dedifferentiation | *Vim* |
| Dedifferentiation | *Runx2* |
| Dedifferentiation | *Fgf18* |
| Dedifferentiation | *Ptk2* |
| Dedifferentiation | *Igfbp3* |

**Table SI 2.** Changes in expression levels and adjusted p-values for genes of interest of PD16 H3WT cells compared to previously published native chondrocyte data shown as normalized count values in Figure SI 6.

| Phenotype | Figure SI 6 Genes | Log2 Fold Change | Log Fold Change Standard Error | Padj |
| --- | --- | --- | --- | --- |
| Chondrogenic | *Sox9* | -3.34 | 0.31 | 6.15 x 10<sup>-27</sup> |
| Chondrogenic | *Col2a1* | -1.64 | 0.41 | 2.55 x 10<sup>-4</sup> |
| Chondrogenic | *Acan* | -5.87 | 0.96 | 7.83 x 10<sup>-9</sup> |
| Dedifferentiation | *Col1a1* | 3.64 | 0.82 | 3.43 x 10<sup>-5</sup> |
| Dedifferentiation | *Thy1* | 5.71 | 1.41 | 1.82 x 10<sup>-4</sup> |
| Dedifferentiation | *Vcan* | -0.24 | 0.43 | 0.662 |
| Dedifferentiation | *Col1a2* | 3.65 | 0.44 | 2.94 x 10<sup>-15</sup> |
| Dedifferentiation | *Col3a1* | 4.33 | 0.33 | 3.50 x 10<sup>-37</sup> |
| Dedifferentiation | *Vim* | 3.39 | 0.22 | 2.00 x 10<sup>-47</sup> |

**Table SI 3.** Changes in expression levels and adjusted p-values for genes of interest of PD16 H3K9M cells compared to previously published native chondrocyte data shown as normalized count values in Figure SI 6.

| Phenotype | Figure SI<br>6 Genes | Log<sub>2</sub> Fold<br>Change | Log Fold<br>Change<br>Standard Error | p<sub>adj</sub> |
| --- | --- | --- | --- | --- |
| Chondrogenic | *Sox9* | -4.01 | 0.25 | 2.86 x 10<sup>-55</sup> |
| Chondrogenic | *Col2a1* | -2.24 | 0.30 | 9.03 x 10<sup>-13</sup> |
| Chondrogenic | *Acan* | -6.12 | 0.40 | 1.10 x 10<sup>-49</sup> |
| Dedifferentiation | *Col1a1* | 3.56 | 0.48 | 1.69 x 10<sup>-12</sup> |
| Dedifferentiation | *Thy1* | 5.79 | 1.05 | 2.19 x 10<sup>-7</sup> |
| Dedifferentiation | *Vcan* | -1.14 | 0.26 | 4.05 x 10<sup>-5</sup> |
| Dedifferentiation | *Col1a2* | 3.59 | 0.39 | 2.53 x 10<sup>-19</sup> |
| Dedifferentiation | *Col3a1* | 4.24 | 0.30 | 6.52 x 10<sup>-44</sup> |
| Dedifferentiation | *Vim* | 3.54 | 0.22 | 9.18 x 10<sup>-53</sup> |

**Table SI 4.** List of p-values for all comparisons in Figure 2 D,E.

| Comparison | Nuclear Area (Fig. 2 Panel D) | Number of H3K9me3 Foci<br>(Fig. 2 Panel E) |
| --- | --- | --- |
| P0 1d to P0 5d | <0.0001 | 0.0038 |
| P0 1d to P0 10d | <0.0001 | 0.0088 |
| P0 1d to P2 1d | <0.0001 | <0.0001 |
| P0 1d to P2 5d | 0.0007 | <0.0001 |
| P0 1d to P2 10d | 0.2535 | 0.1681 |
| P0 1d to P4 1d | <0.0001 | <0.0001 |
| P0 1d to P4 5d | <0.0001 | <0.0001 |
| P0 1d to P4 10d | <0.0001 | <0.0001 |
| P0 5d to P0 10d | 0.0117 | 1.000 |
| P0 5d to P2 1d | <0.0001 | <0.0001 |
| P0 5d to P2 5d | <0.0001 | <0.0001 |
| P0 5d to P2 10d | 0.2287 | 0.9999 |
| P0 5d to P4 1d | <0.0001 | <0.0001 |
| P0 5d to P4 5d | <0.0001 | <0.0001 |
| P0 5d to P4 10d | <0.0001 | <0.0001 |
| P0 10d to P2 1d | <0.0001 | <0.0001 |
| P0 10d to P2 5d | <0.0001 | <0.0001 |
| P0 10d to P2 10d | 0.0007 | 1.0000 |
| P0 10d to P4 1d | <0.0001 | <0.0001 |
| P0 10d to P4 5d | <0.0001 | <0.0001 |
| P0 10d to P4 10d | <0.0001 | <0.0001 |
| P2 1d to P2 5d | 0.0011 | <0.0001 |
| P2 1d to P2 10d | <0.0001 | <0.0001 |
| P2 1d to P4 1d | 0.0535 | 1.0000 |
| P2 1d to P4 5d | 0.0064 | <0.0001 |
| P2 1d to P4 10d | 0.0354 | <0.0001 |
| P2 5d to P2 10d | <0.0001 | <0.0001 |
| P2 5d to P4 1d | <0.0001 | <0.0001 |
| P2 5d to P4 5d | 0.9990 | <0.0001 |
| P2 5d to P4 10d | 0.9960 | <0.0001 |
| P2 10d to P4 1d | <0.0001 | <0.0001 |
| P2 10d to P4 5d | <0.0001 | <0.0001 |
| P2 10d to P4 10d | <0.0001 | <0.0001 |
| P4 1d to P4 5d | <0.0001 | <0.0001 |
| P4 1d to P4 10d | <0.0001 | <0.0001 |
| P4 5d to P4 10d | 1.0000 | 0.9999 |

**Table SI 5.**
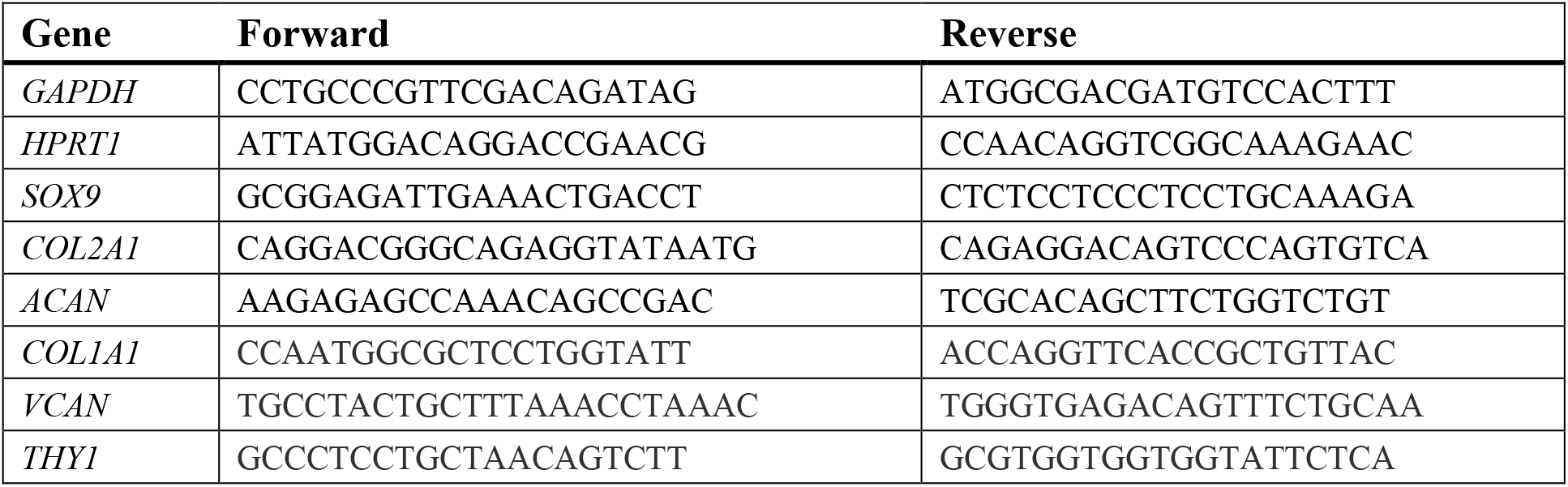
List of chondrogenic and dedifferentiation primers for gene expression analysis using RT-qPCR.

**Table SI 6.**
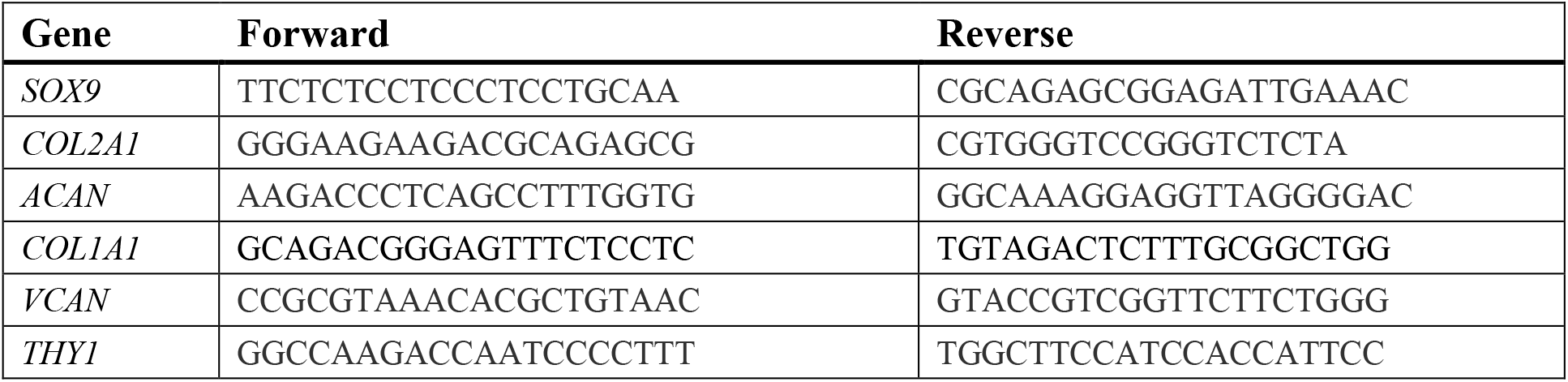
List of qPCR primers for analysis of ChIP isolated DNA.

**Figure SI 1:**
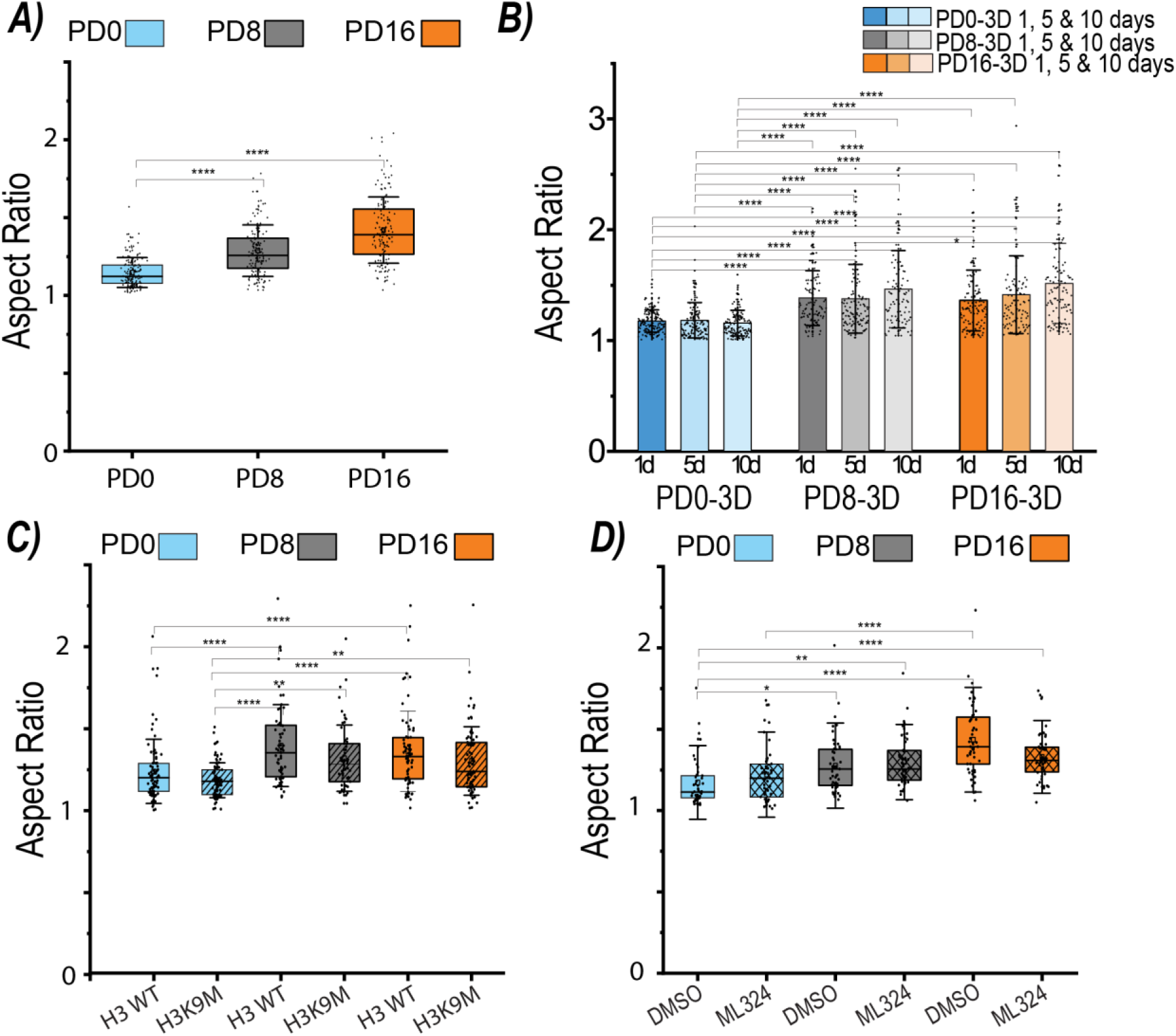
Increased aspect ratio with exposure to a 2D stiff culture was retained in 3D culture and not prevented with suppression of H3K9me3 in 2D culture, but was attenuated with inhibition of KDM4 in 2D culture. Nuclear aspect ratio changes with **A)** increased population doublings, (±s.d.,N=6 biological replicates, n>22 nuclei/treatment/replicate) **B)** time in 3D culture, (±s.d. N=6 biological replicates, n>16 nuclei/treatment/replicate) **C)** suppression of H3K9me3 during the expansion process (± s.d., N=4 biological replicates, n>16 nuclei/treatment/replicate) and **D)** treatment with a KDM4 demethylase inhibitor, ML324 (±s.d. N=3 biological replicates, n>16 nuclei/treatment/replicate). *p<0.05, **p<0.01, ***p<0.001, ****p<0.0001

**Figure SI 2:**
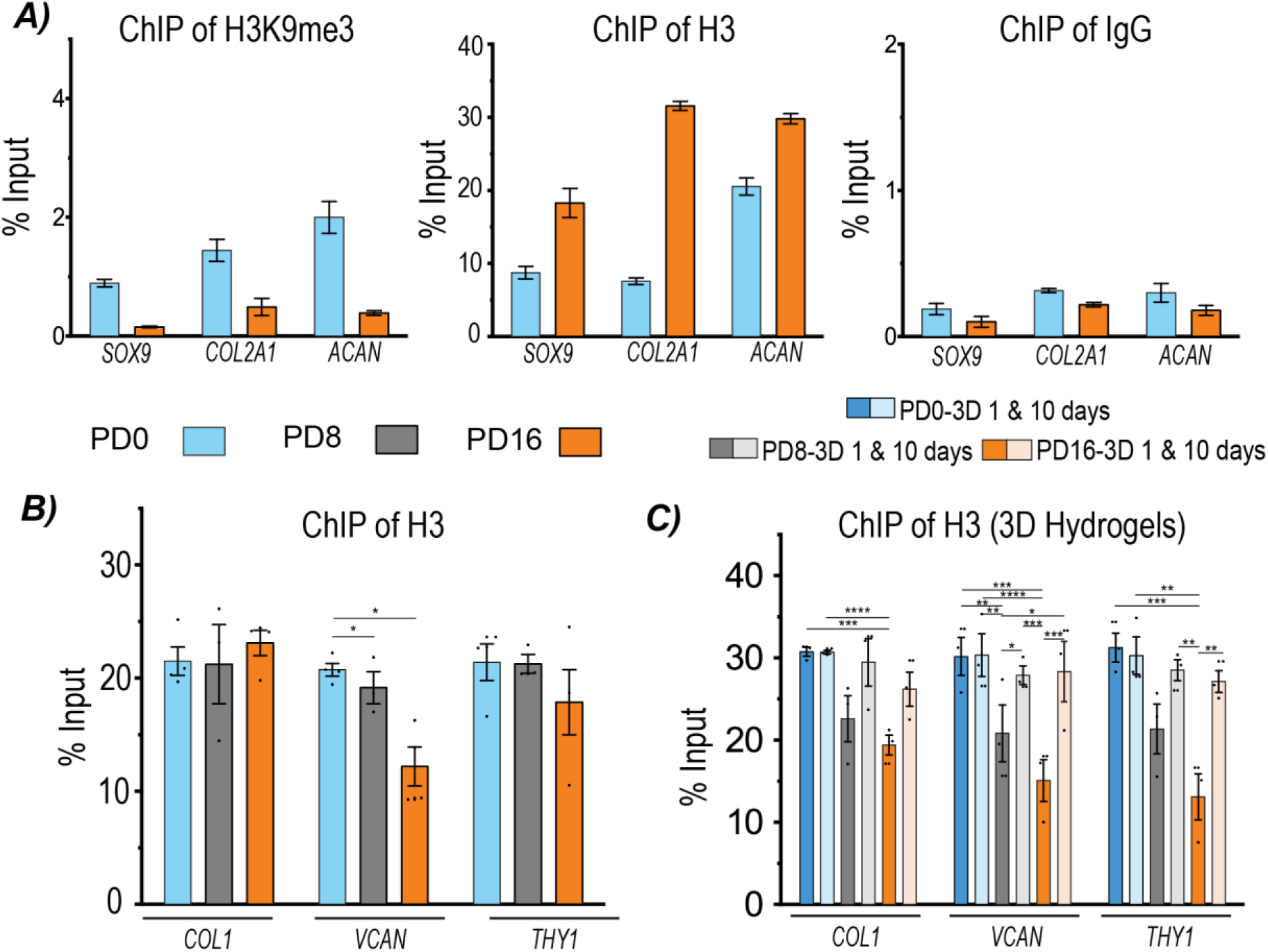
Chromatin immunoprecipitation results did not suggest H3K9me3 regulated the expression of chondrogenic genes, but the measured occupation H3 protein on the promoters of dedifferentiation genes suggested chromatin density influenced expression of dedifferentiation genes in 3D culture. **A)** Chromatin immunoprecipitation results of the occupancy levels of H3K9me3 on the promoters of chondrogenic genes (*SOX9*, *COL2A1*, *ACAN*) (±s.e.m. N=2 biological replicates). **B)** Positive control for ChIP assay, demonstrating the H3 occupancy levels on the promoters of dedifferentiation genes in 2D culture (±s.d. N=3-4 biological replicates). **C)** Positive control for ChIP assay, demonstrating the H3 occupancy levels on the promoters of dedifferentiation genes in 3D culture (±s.d. N=3 biological replicates). *p<0.05, **p<0.01, ***p<0.001, ****p<0.0001

**Figure SI 3:**
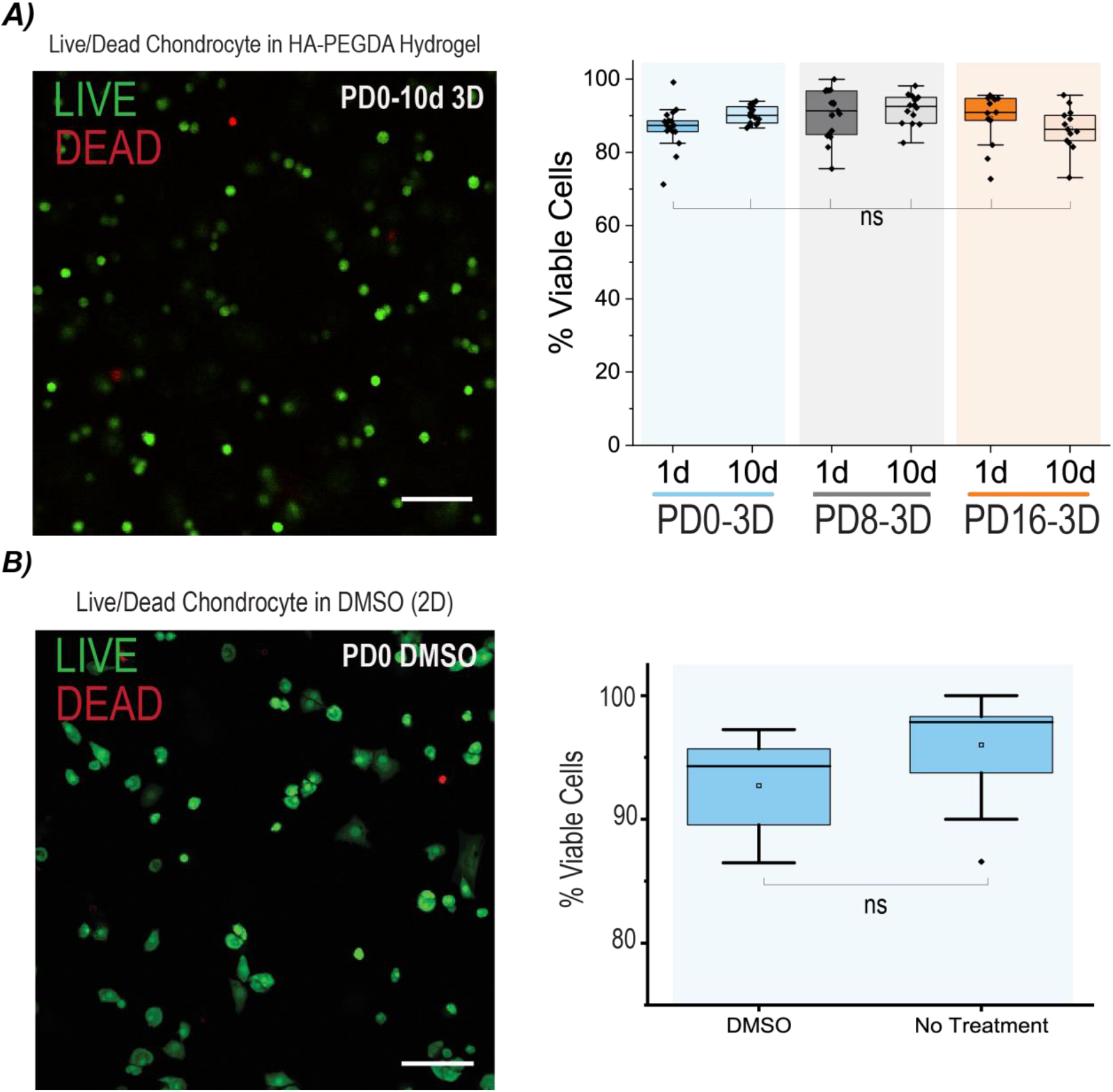
Chondrocytes maintained viability 3D cultures and with DMSO treatment. **A)** Viability of chondrocytes when encapsulated in HA-PEGDA hydrogels and cultured for 10 days with and without prior 2D culture (±s.d. N= 4 biological replicates, n=3-5 images/treatment/replicate). **B)** Viability of chondrocytes cultured with DMSO (±s.d. N= 3 biological replicates, n=3-4 images/treatment/replicate). scale= 100μm, *p<0.05, **p<0.01, ***p<0.001, ****p<0.000, ns=non-significant

**Figure SI 4:**
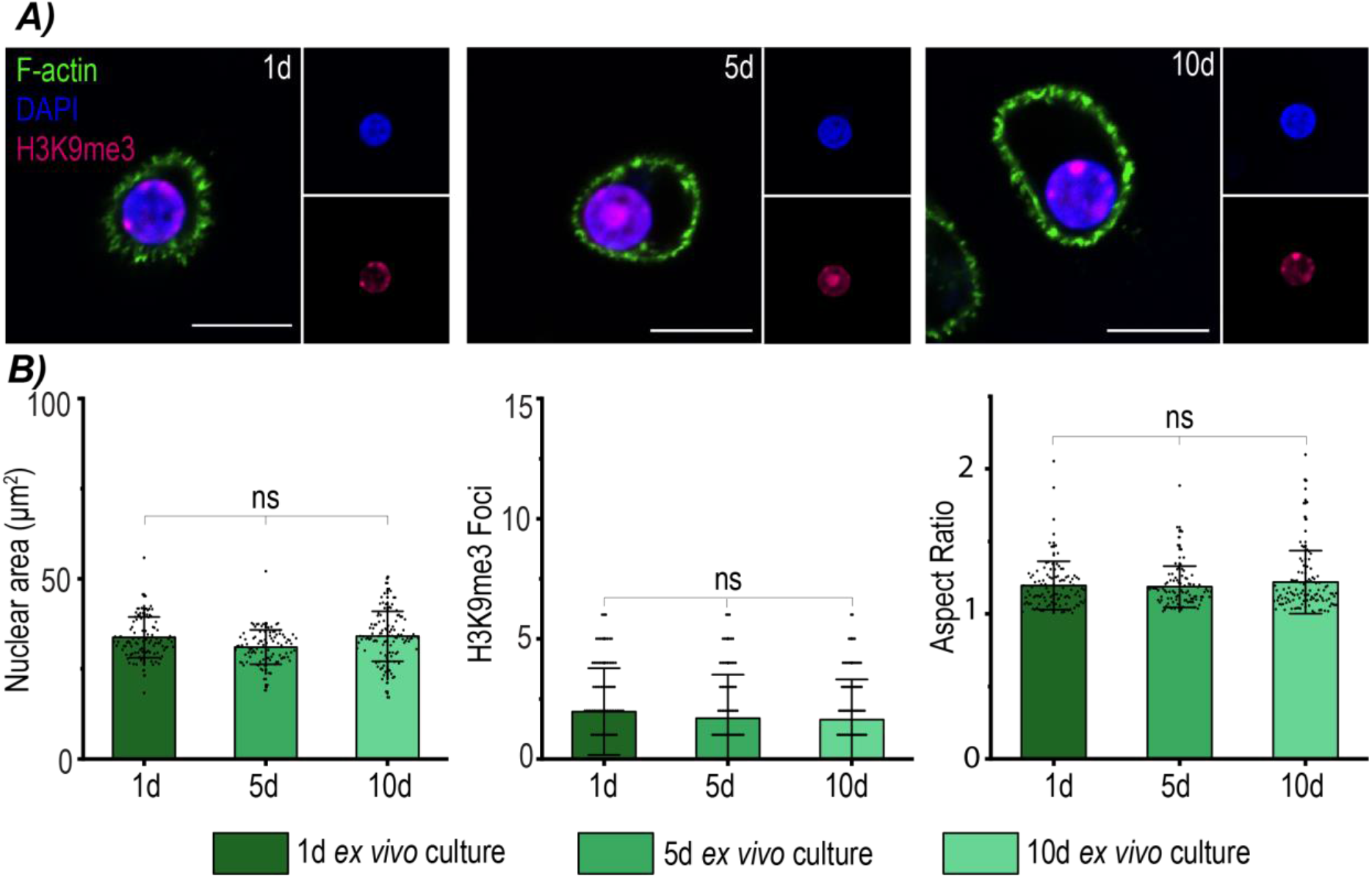
Chondrocytes cultured in the native cartilage environment *ex-vivo* demonstrated similar nuclear architecture characteristics to PD0 chondrocytes encapsulated in 3D hydrogels. **A)** Images of native bovine chondrocytes cultured *ex vivo* for 1, 5 and 10 days (scale= 10μm). **B)** Nuclear areas, nuclear aspect ratio, and H3K9me3 foci of native bovine chondrocytes cultured *ex vivo* for 1, 5, 10 days. (±s.d. N=2 biological replicates, n>20 nuclei/timepoint/replicate). *p<0.05, **p<0.01, ***p<0.001, ****p<0.000, ns=non-significant

**Figure SI 5:**
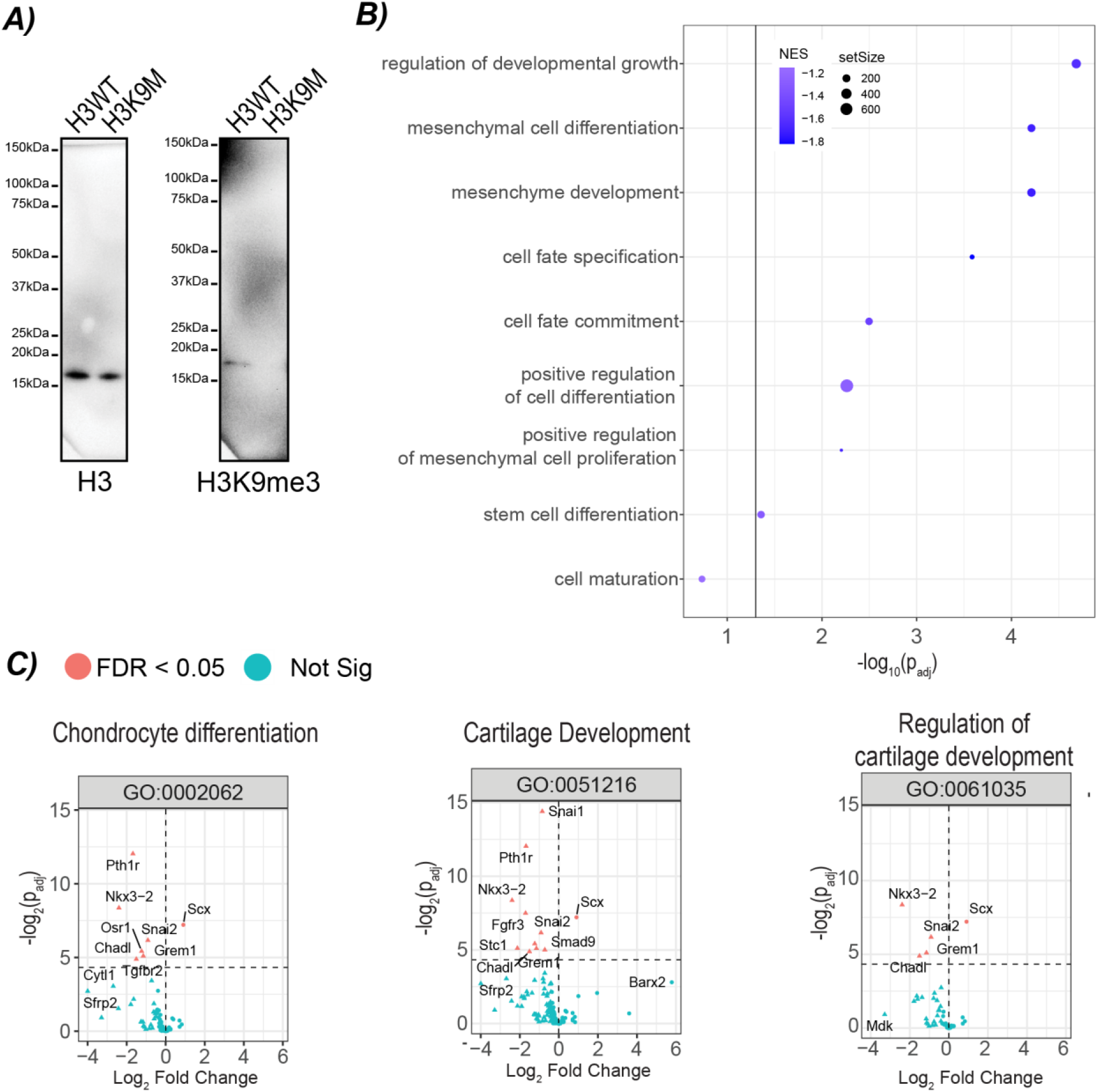
Suppression of H3K9 methylation led to de-enrichment of cell fate and a subset of chondrogenic pathways. **A)** Representative western blot to confirm the suppression of H3K9me3 in comparison to total H3 protein (N=4 biological replicates). **B)** Decrease in H3K9me3 caused suppression of pathways related to cell fate commitment (N=3 biological replicates), and **C)** suppression of genes related to chondrogenic gene ontology pathways (N=3 biological replicates).

**Figure SI 6:**
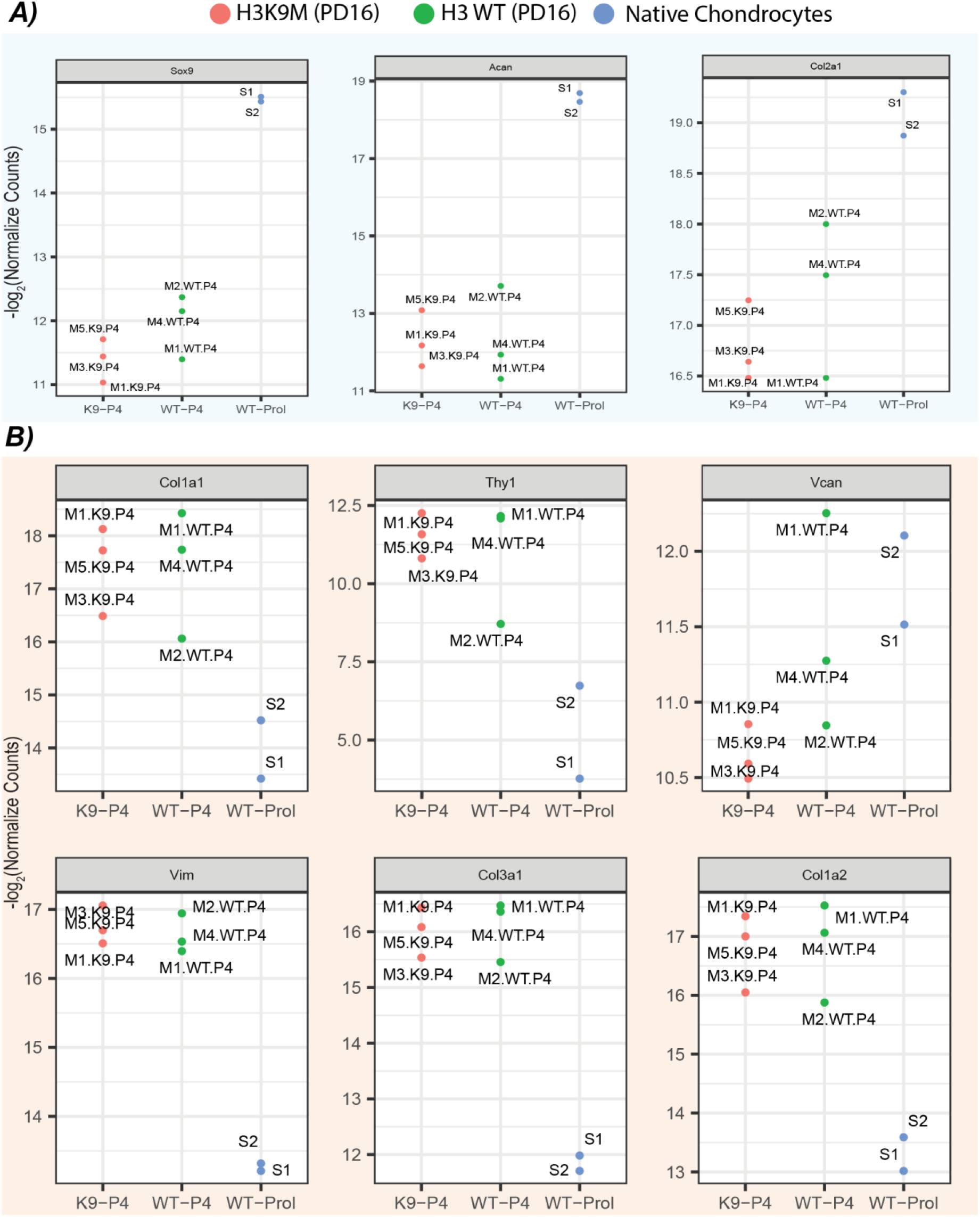
Comparison of RNA-seq data of H3K9M and H3WT cells to published native chondrocyte RNA-seq data, demonstrated that both H3K9M and H3WT chondrocytes dedifferentiated when exposed to 2D stiff substates. Normalized counts of **A)** chondrogenic and **B)** dedifferentiation genes of H3K9M (M1.K9.P4, M3.K9.P4, M5.K9.P4), H3WT (M1.WT.P4, M2.WT.P4, M4.WT.P4), and native (S1, S2) chondrocytes (from previously published data). (N=3 biological replicates for H3K9M/H3WT experiments).

